# Concurrent response and action effect representations across the somatomotor cortices during novel task preparation

**DOI:** 10.1101/2023.10.30.564708

**Authors:** Ana F. Palenciano, Carlos González-García, Jan De Houwer, Baptist Liefooghe, Marcel Brass

## Abstract

Instructions allow us to fulfill novel and complex tasks on the first try. This skill has been linked to preparatory brain signals that encode upcoming demands in advance, facilitating novel performance. To deepen insight into these processes, we explored whether instructions pre-activated task-relevant motoric and perceptual neural states. Critically, we addressed whether these representations anticipated activity patterns guiding overt sensorimotor processing, which could reflect that internally simulating novel tasks facilitates the preparation. To do so, we collected functional magnetic resonance imaging data while female and male participants encoded and implemented novel stimulus-response associations. Participants also completed localizer tasks designed to isolate the neural representations of the mappings-relevant motor responses, perceptual consequences, and stimulus categories. Using canonical template tracking, we identified whether and where these sensorimotor representations were pre-activated. We found that response-related templates were encoded in advance in regions linked with action control, entailing not only the instructed responses but also their somatosensory consequences. This result was particularly robust in primary motor and somatosensory cortices. While, following our predictions, we found a systematic decrease in the irrelevant stimulus templates’ representational strength compared to the relevant ones, this difference was due to below-zero estimates linked to the irrelevant category activity patterns. Overall, our findings reflect that instruction processing relies on the sensorimotor cortices to anticipate motoric and kinesthetic representations of prospective action plans, suggesting the engagement of motor imagery during novel task preparation. More generally, they stress that the somatomotor system could participate with higher-level frontoparietal regions during anticipatory task control.

## 1 Introduction

By using symbolic information, humans convey and communicate task procedures that they have never implemented before. Thanks to this ability, we can acquire novel and arbitrary stimulus-response (S-R) mappings directly from instructions, highlighting the adaptability of human learning processes. This complex skill has been associated with anticipatory brain signals across a set of frontal and parietal areas (Cole et al., 2010; Demanet et al., 2016; Dumontheil et al., 2011; Hartstra et al., 2011, 2012; Palenciano, González-García, Arco, & Ruz, 2019; Ruge & Wolfensteller, 2010) that overlaps with the Multiple Demand Network (MDN; Duncan, 2010; Fedorenko et al., 2013), a system linked to cognitive control. These areas encode in advance behaviorally relevant instruction information, such as the targeted stimuli, the relevant responses, or the rules binding both (Cole, Reynolds, et al., 2013; González-García et al., 2017, 2021; Muhle-Karbe et al., 2017; Palenciano, González-García, Arco, Pessoa, et al., 2019). Accordingly, current models emphasize the role of the MDN in assembling novel task representations from instructions (Brass et al., 2017; Cole, Laurent, et al., 2013). Critically, these representations are characterized by their abstract, multidimensional nature (Bourguignon et al., 2018; Cole et al., 2011; Palenciano, González-García, Arco, Pessoa, et al., 2019). How this higher-order task coding ultimately shapes the downstream sensorimotor processing and impacts instruction implementation remains elusive to date.

To address this issue, we aimed to investigate the contribution of lower-level, sensorimotor neural representations to novel task preparation. In this regard, it has been suggested that internally simulating novel tasks through imagery may facilitate optimal preparation (Moran & O’Shea, 2020). This view resonates with frameworks like the motor simulation theory (Jeannerod, 1994, 2001), which emphasizes that imagery (like planning or other action-related covert processes) is driven by an internal replay that engages action representations shared with overt performance (Grush, 2004; Guillot & Collet, 2005; Hardwick et al., 2018). Hence, by means of imagery or internal simulation, novel instructions could pre-activate sensorimotor neural states overlapping with those driving task execution. Such a mechanism would bridge the above-described abstract task coding found in the MDN with the later behavioral performance. It would also be in line with more general findings from the action preparation domain, where it has been suggested that the motor system anticipates execution-induced neural states during planning (Ariani et al., 2022; Churchland et al., 2010; Shenoy et al., 2013).

Previous behavioral results have evidenced an association between novel task implementation and imagery (Liefooghe et al., 2021; Theeuwes et al., 2018). Robust evidence also emphasizes that just encoding instructions induces automatic response activation before task execution (Liefooghe et al., 2012, 2013; Liefooghe & De Houwer, 2018; Meiran et al., 2015a; Meiran & Cohen-Kdoshay, 2012). Moreover, imposing secondary motor demands during instruction preparation disrupts execution in an effector-specific fashion (Palenciano et al., 2021), suggesting that optimal anticipatory control may rely on task-relevant response representations. At the neural level, functional magnetic resonance imaging (fMRI) data have shown heightened mean activity across motor regions during instruction encoding and preparation (Hartstra et al., 2011, 2012; Ruge & Wolfensteller, 2010). Electroencephalography findings have also evidenced that instructions elicit neurophysiological signatures of response pre-activation (Everaert et al., 2014; Meiran et al., 2015a), namely, lateralized readiness potentials (Gratton et al., 1988). Nonetheless, although all these results converge on the relevance of lower-level motoric processes for novel task preparation, the underlying neural computations have not been systematically explored.

To fill this gap, we addressed whether encoding instructions pre-activated motoric and perceptual neural representations that share content and format with those involved during sensorimotor processing. To do so, we collected fMRI data while participants encoded and executed pairs of novel S-R mappings. Then, they completed a series of localizer tasks seeking to isolate the neural activity patterns of three mappings’ components: the effector-specific motor responses, their somatosensory consequences, and the targeted stimulus categories. Employing canonical template tracking (Palenciano et al., 2023), we estimated standard neural representations (or templates) from the localizer task data and assessed their representational strength during the S-R mapping paradigm. Our main prediction was that the mappings-relevant sensorimotor templates would be anticipatorily encoded in a set of regions of interest (ROIs) located across the motor, somatosensory, and visual brain systems.

## 2 Materials and methods

### 2.1. Participants

Thirty-five participants completed the experiment (27 women, 8 men; mean age = 23 years, *SD* = 3 years). All of them were right-handed, with normal or corrected-to-normal vision, and MRI compatible. Most of the participants were native Dutch speakers (N = 33), except two of them, who were fluent English speakers. All of them signed a consent form approved by the UZ Ghent Ethics Committee. The participation was compensated with 36.25 EURO. The data of four participants were discarded due to excessive movement.

Another participant was unable to finish the MRI session. The data of the remaining 30 participants were included in all the analyses, with the exception of the Action Effects analysis (see Section 2.6), where data from three additional participants needed to be excluded due to technical issues with the tactile stimulators.

### 2.2. Apparatus, stimuli, and protocol

#### Novel S-R mapping paradigm

In the main paradigm, participants encoded pairs of novel S-R mappings to later implement one of them upon target presentation. For that purpose, we generated a pool of unique associations, all consisting of a picture of an animal, and a verbal label indicating the response linked to that stimulus. The mappings’ response involved either the left index, the left middle, the right index, or the right middle finger. The response labels were presented either in Dutch (“*L wijs*”, “*L middel*”, “*R wijs*”, “*R middel*”) or in English (“*L index*”, “*L middle*”, “*R index*”, “*R middle*”) depending on the participant’s preferred language, and were shown in font Arial, size 50 (approx. 1.2° by 3.8°). The pictures were obtained from a previous database containing 770 animate images (Brady et al., 2013; Brodeur et al., 2014; Griffin et al., 2007; Konkle et al., 2010) and completed with stimuli from the Animal Images Database (Possidónio et al., 2019), the C.A.R.E. database (Russo et al., 2018) and available sources with Creative Commons license. All the stimuli were resized to 200 by 200 pixels (approx. 4.8°), and appeared on greyscale, with the background removed and the animal centered on the canvas. Four animal categories were used: canines, felines, birds, and sea animals. For each mapping, we displayed the picture on the left side of the screen, and the response label, on the right side. In each trial, two mappings were shown, one in the upper part of the screen, and another on the lower part.

For each participant, we created a pool of 640 different S-R mappings, arranged in 320 pairs. In each trial, two out of the four possible responses and two out of the four possible animal categories were used. We ensured an equal representation of the six pair-wise responses combination ([left index, left middle], [left index, right index], [left index, right middle], [left middle, right index], [left middle, right middle], and [right index, right middle]) and the six pair-wise combinations of stimulus categories ([cat, dog], [cat, bird], [cat, fish], [dog, bird], [dog, fish], and [bird, fish]). We further controlled the mappings among response and stimulus pairs (i.e.: trials with [cat-left index / dog right-middle] were equally frequent as trials with [bird-left middle / fish-right index]), to avoid that some categories were more frequently linked to some responses by chance. The location of the mappings on the screen (upper or lower part) was also counterbalanced across category and response conditions.

The sequence of events within a trial is depicted in **Fig. 1A**. First, the participants encoded the mappings during 2s, followed by a preparation interval displaying a fixation cross, and with a jittered duration (mean = 2.39s, SD = 1.26s, range = [0.6, 5s], values extracted from a pseudo-logarithmic distribution). Then, a probe stimulus was presented during a window of 1.75s where participants responded. In 90% of the trials, probes corresponded either to the upper or the lower mapping picture (with a 50% probability each). In the remaining 10% of trials (*catch trials*), a novel picture – not in any of the two mappings – was presented, and participants were asked to respond with the left and right index and middle fingers simultaneously. Catch trials were included to ensure that both S-R mappings were equally well encoded. Catch probes were also animal pictures but always belonged to one of the two categories that were not shown in the current mappings. Finally, a fixation was presented during a jittered inter-trial interval (ITI), with the same temporal properties as the previous one.

**Figure 1.**
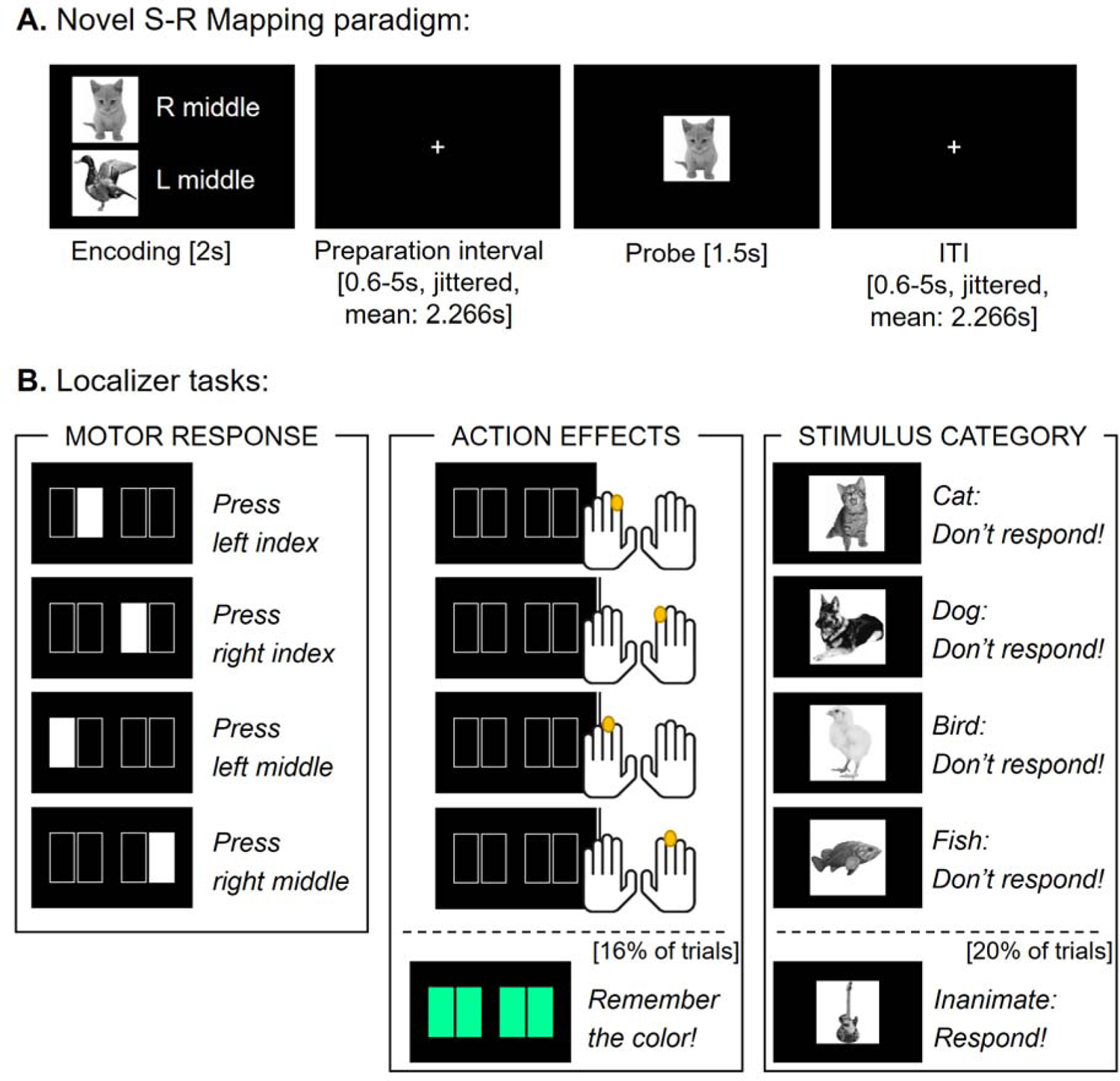
Experimental paradigms. **A.** Trial’s sequence of events during the S-R mapping paradigm. **B.** Localizer tasks. In the motor response localizer (left panel), explicit cues indicated the required response (either with the left or right index or middle finger). In the action effect localizer (middle panel), stimulation was applied separately in each of these four fingers. The visual display consisted of uninformative cues, except for 16% of trials which entailed a secondary, color memory task. In the stimulus category localizer (right panel), pictures of animals belonging to the four categories used in the mappings were interspersed with pictures of inanimate items. Participants responded only to the infrequent (20%) inanimate stimuli.

Before the MRI session, participants practiced the S-R mapping task until they achieved an 85% of accuracy. To ensure that the mappings from the practice session were not further employed, we used a different set of stimuli composed of inanimate items (musical instruments and vehicles). The stimuli from the practice session were not further employed in the experiment. Once inside the MRI scanner, participants completed four task runs, each containing 80 trials. They could take a self-paced pause between runs. Overall, the duration of the S-R mapping task was approximately 45 minutes.

### Localizer tasks

After completing the main paradigm, the participants remained on the scanner and completed three localizer tasks, designed to isolate the representations of the mappings’ motor response, their somatosensory consequences, and stimulus categories. The localizers’ experimental design aimed to maximize the similarity between the targeted component and the S-R mapping paradigm while keeping the remaining task features as independent as possible. This was done to avoid confounding specific template activation with more general motoric or perceptual processes (Palenciano et al., 2023). Each localizer block lasted approximately 5 minutes, and the order in which they were presented was counterbalanced across participants.

In the motor response localizer (Fig. 1B**, left panel**), participants responded to cues that explicitly indicated the required key presses. The cues were composed of four rectangles (200 by 100 pixels, approx. 4.8° by 2.4°) located along the horizontal axis of the screen, each indicating a response with a different finger: from left to right, with the left index, left middle, right index, or right middle finger. The cues were displayed against a black background, and to indicate the required response, the corresponding rectangle was filled in white. We employed these cues instead of verbal labels (as in the S-R mapping task) to avoid semantic processing during the localizer. Instead, we aimed to capitalize on the motor component, by generalizing across instructed response modalities (Liefooghe et al., 2012). Each trial of this localizer consisted of the presentation of the response cue for 1.5s, where participants responded, and a jitter ITI with a fixation cross (same temporal properties as above). Participants completed 80 trials, 20 for each response condition.

In the Action Effect localizer, we aimed to isolate the mappings’ proximal action effects, i.e., the somatosensory consequences of the mapping responses (Fig. 1B**, middle panel**). To do so, we applied single-tap tactile stimulation on the tip of the participants’ left and right index and middle fingers, separately. To do so, we employed four MRI-compatible piezo-electric stimulation devices (mini-PTS, http://www.dancerdesign.co.uk/), each covering an area of 1cm^2^ of the participants’ fingertips. The stimulation pulses were generated with single sawtooth waves, with 40ms of duration. We attached the stimulators to two foam structures, one per hand, in an ergonomic position for the index and middle fingers. The intensity of the stimulation was adjusted to the participants’ sensitivity at the beginning of the MRI session. In the localizer, the stimulation was applied while the cues from the motor response localizer were displayed on the screen. Nonetheless, the cues were not informative about the stimulated finger and all the rectangles appeared filled in black. This was done to keep the two response-related localizers as similar as possible perceptually. Each trial consisted of the pulse of stimulation (40ms) together with the presentation of the uninformative cues for 1.5s, both synchronized at the onset, and a jittered ITI (same temporal properties as above). To facilitate the participants’ engagement during the block, we added a secondary, memory task in 16% of the trials. In these trials, the stimulation was not delivered, and the cues appeared filled in color (pink, yellow, blue, or green). All participants saw the 4 colors in a randomized order, with each color being shown in 4 trials. The participants were instructed to pay attention and to remember the colors displayed, ignoring their appearance order. When the localizer finished, the participants verbally reported the colors that they remembered. The task consisted of 96 trials, and the stimulation was applied in 80 of them (20 trials per stimulation condition).

Finally, participants completed a stimulus category localizer (Fig. 1B**, right panel**), where they discriminated between animal and non-animal items. For each participant, we randomly selected 80 animal pictures (20 per category) and 20 inanimate ones (10 musical instruments, 10 vehicles) from the same database as in the S-R mapping task. All pictures were 200 by 200 pixels, in greyscale, with the background removed and the item centered on the image. In each trial, the stimulus was presented for 1.5s, followed by a jittered ITI (same temporal properties as above). Participants needed to respond simultaneously with their two index and middle fingers only when an inanimate stimulus appeared, which happened in 20% of the trials. In the remaining 80% of trials, an animal stimulus from one of four categories used in the mappings was shown, and no response was required. With this approach, we aimed to isolate the perceptual processing of the four animal categories, in the absence of any motor preparation or execution. The localizer consisted of 100 trials, 80 displaying an animal picture (20 per animal category).

All tasks were programmed in Psychopy (v3; Peirce et al., 2019) and were presented on a screen located at the back of the scanner and projected onto a mirror system. Responses were collected using a pair of MRI-compatible joysticks.

### 2.3. Experimental design

In the S-R mapping paradigm, we manipulated two within-subjects variables: the mappings’ relevant responses ([left index, left middle], [left index, right index], [left index, right middle], [left middle, right index], [left middle, right middle], and [right index, right middle]), and stimulus category ([cat, dog], [cat, bird], [cat, fish], [dog, bird], [dog, fish], and [bird, fish]). We collected 48 trials per response and per category condition. For experimental control, we fully crossed these two variables, resulting in 36 different mappings pairs, sampled through eight observations each. However, all our analyses treated independently the response and category manipulations (e.g.: in the response-related analyses, we collapsed across stimulus categories within response conditions). Hence, our analyses were based on 48 observations per cell.

In the motor response and action effects localizers, we had one within-subject manipulation: the effector (left index, left middle, right index, and right middle) used to respond or stimulate. In the stimulus category localizer, we manipulated within subjects the stimulus animacy (animate, and inanimate items) and the animal category of the animate stimulus (cat, dog, bird, and fish). In the three localizers, we collected 20 observations per condition of interest.

### 2.4. MRI data acquisition and preprocessing

The MRI data was collected with a 3T Siemens Magnetom Trio scanner at the Ghent University Hospital (UZ Ghent, Belgium) with a 64-channel head coil. Whole-brain functional images were obtained with a T2* EPI sequence [TR = 1730 ms, TE = 30 ms, flip angle=66°, FOV = 210 mm, image matrix = 84 × 84, voxel size = 2.5 x 2.5 x 2.5 mm, distance factor = 0%, 50 slices, slice acceleration factor = 2, slice orientation according to the participants’ AC-PC line]. Four 400-volumes runs were collected for the S-R mapping task, and three 185-volumes runs for the localizers. Before the functional runs, we collected anatomical T1-weighted images using a MP-RAGE sequence [TR = 2250 ms, TE = 4.18 ms, flip angle=9°, FOV = 256 mm, image matrix = 256 x 256, voxel size = 1 x 1 x 1 mm, 176 slices]. We also collected phase and magnitude field map images [TR = 520 ms, TE1 = 4.92 ms, TE2 = 7.38 ms, flip angle=60°, FOV = 210 mm, image matrix = 70 × 70, voxel size = 3 × 3 × 2.5 mm, distance factor=0%, 50 slices], to correct for inhomogeneities of the magnetic field. Overall, participants remained inside the scanner for 75 minutes.

MRI Data preprocessing and GLM estimation (see Section 2.6) were carried out on SPM12 (http://www.fil.ion.ucl.ac.uk/spm/software/spm12/). The functional images were spatially realigned, corrected for magnetic field inhomogeneities, and slice-time corrected. The T1 anatomical images were defaced, coregistered to the participants’ mean functional volume, and segmented. For univariate analyses, the functional images were normalized to the MNI space using the forward deformation fields derived from the T1 segmentation and smoothed with an 8mm FWHM Gaussian kernel. The multivariate analyses were performed on unnormalized and unsmoothed images to avoid distorting the finer-grained activity patterns. The normalization and smoothing were performed on the results maps. The T1 segmentation also generated inverse deformation fields which were used to convert the ROIs masks, defined in the MNI space, into the participants’ native anatomical space.

Four participants showed sustained, excessive movement (>2.5mm) and were removed from the sample. However, 3 additional participants displayed punctual movements larger than our voxel size but restricted on time. For those participants, the volumes affected by movement (less than 1% of the acquired images) were masked in further analyses using regressors generated with the ART toolbox (http://web.mit.edu/swg/software.htm).

### 2.5. Statistical analyses: behavioral data

We analyzed the effect of the mappings’ relevant responses and stimulus categories on response accuracy and speed, separately. This was done in an exploratory fashion, to assess whether the conditions were behaviorally balanced. We carried out repeated-measures ANOVAs, using as within-subject factor either the response or the category condition. A Greenhouse-Geisser correction was used when sphericity was violated. Post hoc pair-wise *t*-tests were carried out to identify conditions where performance deviated from the rest. A Bonferroni-Holm correction for multiple comparisons was applied. Catch trials were excluded from the analyses. Only correct trials were included in RT ANOVAs.

### 2.6. Statistical analyses: fMRI data

#### Univariate analyses

We first conducted a series of univariate analyses to provide an overview of the regions engaged by both the S-R mapping paradigm and the localizer tasks. For the S-R mapping task, we fitted a GLM with regressors covering the mapping encoding stage, the preparation interval, and the probe, separately for the six response conditions. Each regressor was defined using the event duration, except the probe, which was modeled with the trial RT. The ITI was not defined in the model and thus contributed to the implicit baseline. As nuisance regressors, we included errors, catch trials, and the six movement parameters derived from the realignment. Within participants, we carried out three one-sample *t*-tests to contrast each event of interest (encoding, preparation, and probe) against the baseline, collapsing across conditions. The outputs were entered into group-level *t*-tests to detect significant BOLD signal increases induced by our paradigm’s events. Both the GLMs and the individual and group-level *t*-tests were estimated on a voxel-by-voxel basis, following the traditional univariate approach. The results were later corrected at *p* < .05 using a cluster-wise FWE multiple-comparison correction (from an initial, uncorrected threshold of *p* < .001 and k = 10). For all the whole-brain analyses, the T values reported correspond to the cluster’s peak.

The three localizers were analyzed in separate GLMs. For the motor response localizer, we defined a regressor for each response condition, locked to the onset of the response cue and with duration fixed to the trials’ RT. Errors were included as a nuisance regressor. For the action effects localizer, the GLM included a regressor for each stimulation condition, locked and with the same duration as the stimulation pulse. Trials from the secondary memory task (i.e., when colored cues appeared, and no stimulation was delivered) were included as a nuisance regressor. Finally, for the stimulus category localizer, we modeled a regressor per animal category, with the same onset and duration as the stimulus presentation. Inanimate target trials and errors were included as nuisance regressors. The localizers GLMs also included the six motion parameters. We computed one-sample *t-*tests to detect BOLD signal increases linked to the localizers’ event of interest (response, tactile stimulation, and target presentation) against the implicit baseline. We followed the same strategy as with the S-R mapping task regarding voxel-level GLM and *t*-tests estimation and cluster-wise FWE-based statistical inference.

#### Canonical template tracking

The main goal of this study was to assess whether the motoric and visual representations guiding overt performance and perception were covertly engaged during novel mapping preparation. To do so, we focused on patterns of brain activity distributed across neighboring voxels, both within a series of ROIs and using an unconstraint whole-brain procedure (see below). These activity patterns were analyzed using canonical template tracking, a multivariate technique that enables the comparison of neural data between a targeted cognitive paradigm and theoretically driven localizer tasks (Palenciano et al., 2023; for other examples, see González-García et al., 2021; Wimber et al., 2015). Specifically, we estimated a series of canonical neural representations (or templates) from each localizer data. These templates aimed to capture the corresponding motoric or perceptual component representation in the same format as engaged during overt processing (i.e., pressing a key, experiencing the action effect, or perceiving the stimulus category). Then, we assess the representational strength of these templates during the S-R mapping paradigm, by computing their similarity with the multivoxel activity patterns evoked by the novel mappings’ preparation interval.

First, we fitted a set of GLMs to estimate the multivoxel activity patterns during the S-R mapping task (Fig. 2A). To maximize the number of data points being fed into the template tracking procedure, we estimated the signal in a trial-by-trial fashion. We used a least-square-sum (LSS) approach (Arco et al., 2018; Mumford et al., 2012, 2014) to reduce the collinearity. For each trial, we estimated a separate GLM with the following regressors: (1) the preparation interval of the current trial; (2) the encoding stage, collapsing across all correct trials; (3) the preparation stage, excluding the current trial but collapsing across the remaining correct trials; (4) the probe event, collapsing across all correct trials. All trial-wise GLMs further included nuisance regressors modelling, (5) errors and (6) catch trials (using the same onset and duration as in the univariate GLM defined above), and the six movement parameters. We iterated the estimation procedure across all the trials, obtaining a beta map for the preparation interval of each pair of novel mappings. Next, we performed one-sample *t*-tests comparing each beta map against the baseline, to normalize noise from a univariate perspective (Kriegeskorte et al., 2008; Walther et al., 2016). The activity patterns used for the template tracking were directly extracted from the resulting trial-wise *t*-maps.

**Figure 2.**
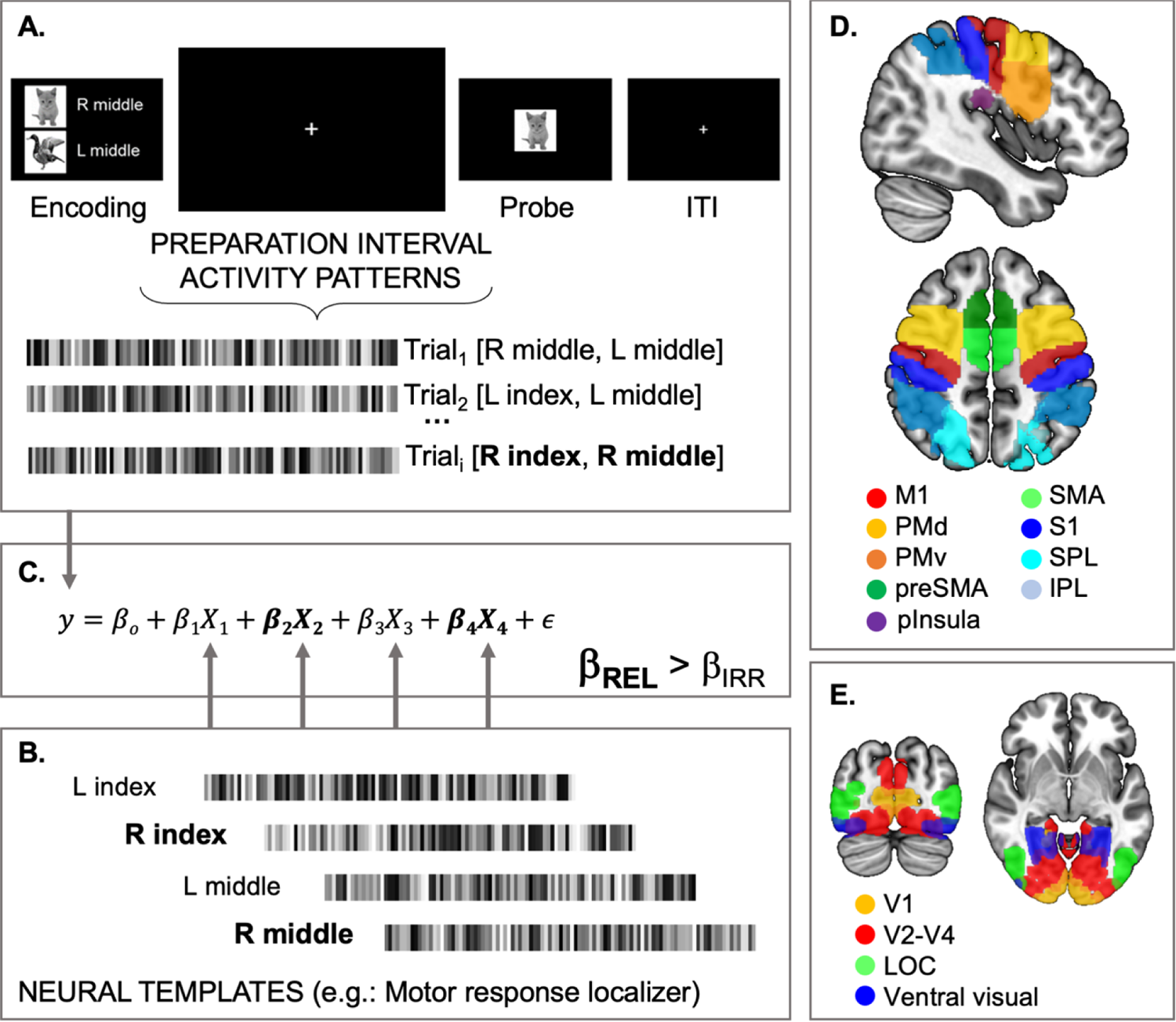
Canonical template tracking. **A.** Estimation of trial-by-trial activity patterns from the preparation interval of the S-R mapping paradigm. In the trial shown as an example (Trial_i_), the prepared mapping responses entailed the right index and middle fingers. **B.** Canonical template estimation. From each localizer, we estimated a single activity pattern per condition. Here we illustrate the motor response localizer, with templates corresponding to the four response conditions. The templates relevant to the mapping are shown in bold. **C.** A multiple regression was fitted to predict each trial’s activity pattern. The regressors entailed the four (two relevant, in bold, and two irrelevant) canonical templates, together with a constant and an error term. The inference was carried out on the comparison between averaged beta weights from relevant and irrelevant templates. **D.** ROIs used in the motor response and action effect analyses. **E.** ROIs used in the stimulus category analyses.

Next, we estimated the canonical neural templates using the three localizers’ data (Fig. 2B). To maximize the robustness of the templates, we estimated standard GLMs that provided a single beta map per localizer condition. Within each localizer, the four conditions’ beta images (corresponding to the four effectors in the case of the motor response and action effects localizers, and to the four animal categories in the stimulus category localizer) were contrasted separately against the baseline. The templates were extracted from the resulting *t*-maps.

Finally, we estimated the similarity between the templates and each activity pattern from the mappings’ preparation interval (Fig. 2C). In each trial from the S-R mapping task, two out of the four possible motor responses, and two out of the four possible stimulus categories, were prepared in advance. This design allowed distinguishing between two subsets of templates regarding a given trial: relevant (i.e., prepared) and irrelevant (i.e., not prepared) ones. Hence, we estimated the similarity between the main task data and both types of templates and used the irrelevant similarity measurements as baseline. Critically, the use of this baseline condition allowed us to infer the specificity of the motoric and perceptual information encoded during the preparation (Palenciano et al., 2023). To compare the similarity estimates from several templates simultaneously, while accounting for the shared variance among them, we used a multiple regression framework (e.g., Palenciano et al., 2019).

We first carried out the template tracking separately for each localizer task and on a set of ROIs (see *ROI selection* section). For each region, we first normalized (i.e., z-scored) each multivoxel activity pattern from the S-R mapping task and each canonical template, ensuring that, across voxels, the mean activity was 0 and the standard deviation was 1. That sought to mitigate the impact of univariate activity differences on our canonical template results. Then, we fitted a series of multiple regressions in which the multivoxel activity pattern from a trial’s preparation interval was predicted using the four localizer templates as regressors. That provided beta coefficients that reflected the similarity between relevant and irrelevant templates and the main task data. Then, we averaged the beta weights across templates and trials to obtain a single relevant and irrelevant value per ROI. At the group level, we carried out one-tailed, paired-sample Wilcoxon signed rank tests to assess the hypothesis that the beta weights from relevant templates were higher than the irrelevant ones. This comparison offered an estimate of the representational strength of specific templates while controlling for more general motor preparation or visual expectation processes. Furthermore, one-sample Wilcoxon tests were performed to assess whether relevant and irrelevant beta weights were different from zero. It is important to note that the later results could be contaminated by the positive bias present in similarity estimates, and hence, were only interpreted to qualify the comparison against the irrelevant (or baseline) condition. In all cases, we controlled for multiple comparisons (i.e., number of ROIs) using a Bonferroni-Holm correction (Holm, 1979).

Since the motor response and the action effects localizers tapped into a common component of the S-R mappings, we performed an additional analysis including templates from the two localizers simultaneously. With this approach, we aimed to disentangle the contribution of passive perception of the action effects, from a richer action representation conveying motor planning and execution. The procedure was identical to the one specified above, with the exception that the multiple regressions included eight regressors (one per effector condition and localizer) instead of four. After averaging, we obtained four estimates per ROI, corresponding to relevant and irrelevant motor response and action effects templates. Next, we carried out a repeated measures ANOVA including the ROI, the localizer task, and the template condition (relevant, irrelevant) as within-subject factors.

We carried out post hoc comparisons to explore the three-way interaction between ROI, Localizer, and Template, using a Bonferroni-Holm multiple comparison correction.

#### Response templates: exploratory analyses

In addition to our planned analyses, we aimed to better understand which aspects of the mappings’ instructed responses were captured by our canonical templates. In this regard, the novel mappings involved different response effectors (index, middle fingers) and different laterality (left-hand, right-hand responses). Due to the pervasive effects of response laterality, we aimed to address whether specific response effector representations could be also detected with the canonical template procedure. While the experiment was designed to control for laterality, by balancing it across the response conditions, we further investigated this issue by repeating our main analysis in a subset of novel mappings where only the templates’ effector information (and not the laterality) was relevant.

First, we extracted the relevant and irrelevant beta weights from S-R mappings that entailed the same effector from the two hands (i.e., the conditions [left index, right index] and [left middle, right middle]) and estimated the activation of the relevant effector templates in comparison with the irrelevant ones. In these two targeted conditions, the relevant and irrelevant templates would differ only regarding the response effector, being equated regarding response laterality. For example, in the [left index, right index] condition, the relevant templates encompassed the two (left and right) index fingers, while the irrelevant ones consisted of the two (left and right) middle fingers, so that the comparison between both only entailed different effectors (index vs. middle fingers) but not literalities. That would not be the case of the conditions excluded from this test: for instance, in the [left index, left middle] condition, the two left-hand relevant templates would be compared against the two right-hand irrelevant ones. This analysis was carried out in all the ROIs examined in the main analysis.

Second, to provide a deeper insight into the processes implemented in the primary motor and somatosensory cortex, we carried out an additional test on these ROIs using the same subset of trials but further differentiating the laterality of the templates. Specifically, each canonical template was estimated using data from the contralateral hemisphere ROIs. For instance, to predict the activity patterns of mappings involving [left index, right index], we used data from the left M1 to estimate the similarity with the right index (relevant) and right middle (irrelevant) templates, and data from the right M1 to estimate the similarity with the left index (relevant) and left middle (irrelevant) templates. Hence, this test replicated the above-described logic but considering the contralateral functional organization of the primary somatomotor cortices. This way, we ensured that our results were driven by specific effector representations engaged in the corresponding, contralateral primary cortex. Nonetheless, it is important to stress the post hoc nature of these two analyses, for which the experiment design was not optimized. As a result, only 33% of the observations from the S-R mapping task were included, with a corresponding negative impact on the analysis’s statistical power. Still, these results helped qualify the planned analyses here reported.

Finally, we aimed to ensure that our main findings were not driven by motoric brain signals that spread from the probe epoch into the preparation interval. Considering the temporal proximity of these two events and the sluggishness of the fMRI BOLD signal, it could be argued that signals originated during task execution could contaminate the preparatory activity patterns. To prevent this confound, we employed a validated jittered distribution (e.g., González-García et al., 2021) and used an LSS estimation protocol that facilitated signal segregation (Arco et al., 2018; Mumford et al., 2012, 2014). Still, we performed a series of further tests to evidence that template instantiation could be detected in the absence of later action. In our task, only one of the two S-R mappings – and thus, prepared responses –were actually executed after the preparation interval. Hence, we focused on the prepared but not executed mappings and repeated the above-described response-related canonical template analyses. Specifically, we replicated the ROI-based multiple regression (using the motor response and action effects templates both independently and simultaneously) including a single relevant template: the one corresponding to the prepared response that was not executed in the current trial. Replicating the same pattern of results in these analyses would strongly argue against signal misattribution.

#### ROI selection

We used two sets of a-priori ROIs, one for the motor response and action effects templates, and another one for the stimulus category ones. In the former case, we selected regions associated with action control and somatosensory processing. Specifically, we focused on the primary motor (M1) and somatosensory cortex (S1), the ventral and dorsal portion of the premotor cortex (PMv and PMd, respectively), the supplementary and pre-supplementary motor areas (SMA and preSMA, respectively), and the inferior and superior parietal lobe (IPL and SPL, respectively). A post hoc ROI was placed on the posterior insula (pInsula) due to its involvement during the tactile stimulation processing (see Fig. 4). The impact of the stimulus category templates was assessed in regions processing both physical stimulus attributes and more abstract, categorical information. Specifically, we used as ROI the primary visual cortex (V1), the early visual cortices from V2 to V4 (V2-V4), the lateral occipital complex (LOC), and a broader region covering the ventral pathway until the inferotemporal cortex (VentralVisual).

The M1, PMv, PMd, SMA, preSMA, and S1 ROIs were obtained from the Human Motor Area Template (HMAT; Mayka et al., 2006), while the IPL, SPL, V1, V2-V4, LOC, and VentralVisual masks were extracted from the HCP-MMP1.0 parcellation (Glasser et al., 2016). All ROIs covered the bilateral portions of the corresponding region, except for the M1 and S1, for which we used separated ROIs for the left and right hemispheres to account for their contralateral functional organization. This was done to avoid capturing broader interhemispheric differences in univariate signal that would remain after the z-score correction. These ROI masks comprised a range from 2120 to 15489 voxels in the standard MNI space (left M1: 3761 voxels; right M1: 3761 voxels; left S1: 2120 voxels; right S1: 2144 voxels; PMv: 5809 voxels; PMd: 6607 voxels; SMA: 2722 voxels; preSMA: 2206 voxels; IPL: 3743 voxels; SPL: 4265 voxels; V1: 11184 voxels; V2-V4: 24841 voxels; LOC: 9427 voxels, VentralVisual: 15489 voxels). Finally, we used MarsBar to generate the pInsula ROI, building two 10mm radius spheres located in the bilateral activation peaks found in the univariate results from the action effect localizer. The bilateral pInsula mask comprised 1030 voxels.

For each localizer, we included a control ROI that was unrelated to the templates of interest and where no effects were expected. Specifically, we used V1 in the motor response and action effects analyses, and M1 in the stimulus-related analysis.

#### Whole-brain searchlight analyses

To complement the ROI results, we additionally performed exploratory whole-brain analyses using a searchlight procedure. We iterated a 5-voxel radius sphere across every location of the brain, performing multiple regressions with the S-R mapping task activity patterns and templates estimated from the current sphere data. The averaged beta coefficients for the relevant and irrelevant templates were assigned to the voxel at the center of the sphere. As a result, we obtained two beta maps per participant and localizer task, one for the relevant and another for the irrelevant templates. Next, within participants, we subtracted the irrelevant beta map from the relevant one. The resulting images were normalized and smoothed, and entered into group-level one-sided, one-sample *t*-tests to detect significantly greater beta weights linked to relevant templates, in comparison with irrelevant ones.

We further used the searchlight whole-brain maps to formally test whether univariate and multivariate preparatory signals overlapped anatomically. To do so, we thresholded (*p* < .05, FEW corrected) and binarized the group-level brain maps obtained in the univariate *t*-test from the preparation interval and the three canonical template tracking searchlights. Then, we computed the conjunction between each pair of univariate and multivariate maps to identify the voxels that remained significant in the two contrasts (Nichols et al., 2005).

#### Control analysis: template reliability

Finally, we aimed to ensure that our results were not driven by differences in the reliability of the canonical templates employed. The experimental design sought to eliminate such confound by balancing each localizer’s set of templates (e.g., the four response effectors’ templates) between the relevant and irrelevant conditions. Nonetheless, we computed two reliability measurements per template (González-García et al., 2021; Palenciano et al., 2023). First, to estimate the overall univariate noise, we computed the Signal-to-Noise Ratio (SNR), dividing the absolute mean t value from a given template by its standard deviation. Then, to assess the stability of the templates’ informational content, we used a correlationability index (González-García et al., 2021). We split each localizer data into odd and even trials and fitted new GLMs, obtaining two (instead of a single) canonical templates per localizer condition. The correlationability index consisted of the Pearson correlation coefficient between each condition’s pair of templates. To identify reliability differences among templates, we run one-way repeated measures ANOVAs separately for each measurement, localizer, and ROI. The template condition was used as a single factor. Post hoc paired-sample *t*-tests (Bonferroni-corrected for multiple comparisons) were computed whenever the main effect of the template condition was significant.

## 3. Results

### 3.1. Behavioral results

Performance was in line with previous studies using novel S-R associations. On average, participants responded accurately on 89% of trials of the S-R mapping task (SD = 6%), with a mean RT of 710 ms (SD = 99 ms). On catch trials, the average accuracy was 84% (SD = 22%), and the mean RT, 943 ms (SD = 134 ms). Regarding the localizers’ performance, in the motor response task, participants responded correctly in 98% of trials (SD = 1.7%), with an average RT of 523 ms (SD = 58 ms). In the action effects localizer, all the participants recalled correctly the four colors used in the secondary memory task. Finally, in the stimulus category localizer, the mean accuracy was 99% (SD = 12%), with an average of 0.2% of false alarms (SD = 0.5%) and 4% of misses (SD = 8%). The average RT was 942 ms (SD = 14 ms).

We explored whether the behavioral performance was modulated by the mappings’ relevant responses and stimulus categories. The response condition effect (Fig. 3) was not significant neither accuracy rates, F(3.4, 99.9) = 0.83, *p* = .49, η_p_^2^ < .01, nor in RTs, F(5, 145) = 1.36, *p* = .24, η_p_^2^ = .05, supporting that performance was equivalent across response pairings. Regarding the stimulus category (Fig. 3), however, we found a significant effect on both accuracy rates, F(5, 145) = 3.84, *p* = .003, η_p_^2^ = .12, and RTs, F(5, 145) = 2.38, *p* = .04, η_p_^2^ = .08. Post hoc paired-sample *t*-test showed that, in general, participants performed worst mappings that included the category pair [bird-fish]. Accuracy was lower in this condition than in the [cat-dog] pair (T = 3.43, *p* = .01) and the [cat-fish] pair (T = 3.90, *p* = .002), with an additional marginal difference with the [cat-bird] pair (T = 2.84, *p* = 0.07). RTs were marginally slower in [bird-fish] trials than in [dog-fish] ones (T = 2.79, *p* = 0.09). More qualitatively, the overall pattern reflected a general impoverishment in the [bird-fish] category pairings than in the other conditions.

**Figure 3.**
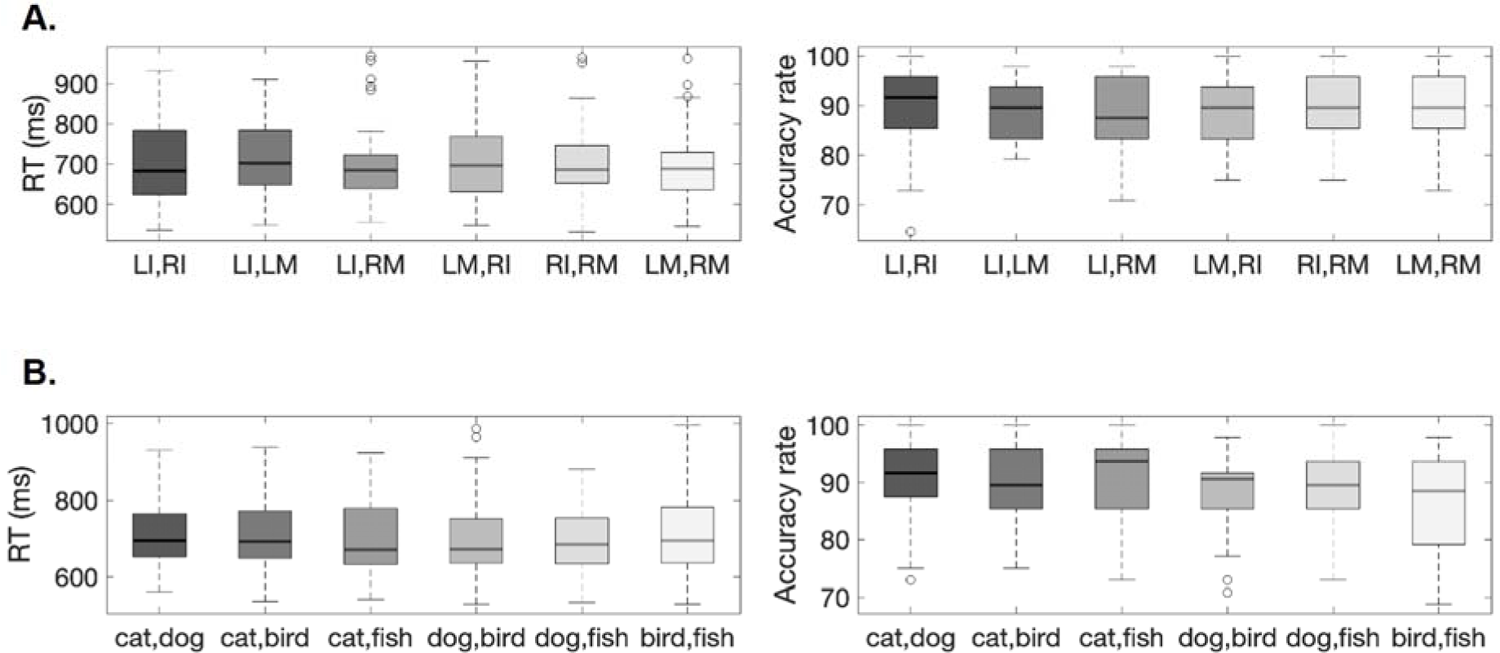
Boxplots displaying the RTs (left) and percentage of correct responses (right) during the novel S-R Mapping paradigm across the six response conditions **(A)** and six stimulus category conditions **(B)**. LI = left index; RI = right index; LM = left middle; RM = right middle.

Finally, to explore whether the S-R mappings combination further modulated behavioral performance we carried out additional repeated measures ANOVA including the response and the category condition as factors. In line with the above-described results, the effect of category was significant for both RT, F(4.31, 124.84) = 2.54, *p* = .03, η_p_^2^ = .08, and accuracy rates, F(4.22, 122.23) = 3.84, *p* = .003, η_p_^2^ = .12. Contrary, neither the response condition nor the interaction was significant (all *p values* > .20).

### 3.2. fMRI results

#### Univariate analyses

As a first step, we aimed to characterize the regions engaged by the different stages of the S-R mapping paradigm. Replicating previous findings, encoding novel S-R associations was associated with increased activity broadly in the MDN and the visual and motor systems (Fig. 4A). A wide cluster ([38, −66, −16], k = 57033, T(1,29) = 20.05, *p* < .001) covered the early and late visual cortices, extending dorso-medially into the precuneus and anteriorly into the inferotemporal cortex. The cluster also included S1, M1, PMC, and SMA. Within the MDN, it covered the lateral DLPFC, incurring into the anterior insula/frontal operculum (aI/fO) in the left hemisphere, and the SPL, IPL, and IPS. It also covered part of the cerebellum and the thalamus. A second cluster ([32, 22, 4], k = 539, T(1,29) = 7.50, *p* < .001) covered the right aI/fO. Conversely, during the mappings’ preparation interval (Fig. 4B), only a restricted set of regions showed a consistent activity increase. Specifically, we found significant clusters bilaterally in the aI/fO (left: [−30, 20, −2], k = 867, T(1,29) = 13.49, *p* < .001; right: [34, 22, −2], k = 825, T(1,29) = 14.38, *p* < .001), the inferior frontal junction (left: [-42, 24, 28], k = 258, T(1,29) = 5.58, *p* = .001); right: [40, 30, 30], k = 442, T(1,29) = 7.28, *p* < .001), the preSMA ([6, 20, 50], k = 1323, T(1,29) = 11.61, *p* < .001), and the bilateral IPS and IPL (left: [−32, −54, 42], k = 376, T(1,29) = 6.60, *p* < .001; right: [36, −54, 42], k = 387, T(1,29) = 7.15, *p* < .001). Finally, probe processing (Fig. 4C) substantially engaged the visual and motor system and the MDN, as well as subcortical nuclei. An extended cluster ([36, 2, 10], k = 104250, T(1,29) = 19.61, *p* < .001) overlapped with the activations found during the encoding, also including the premotor cortices, the preSMA, the middle cingulum, the middle and posterior region of the insula, the temporal operculum, the rostrolateral prefrontal cortex, and the basal ganglia.

**Figure 4.**
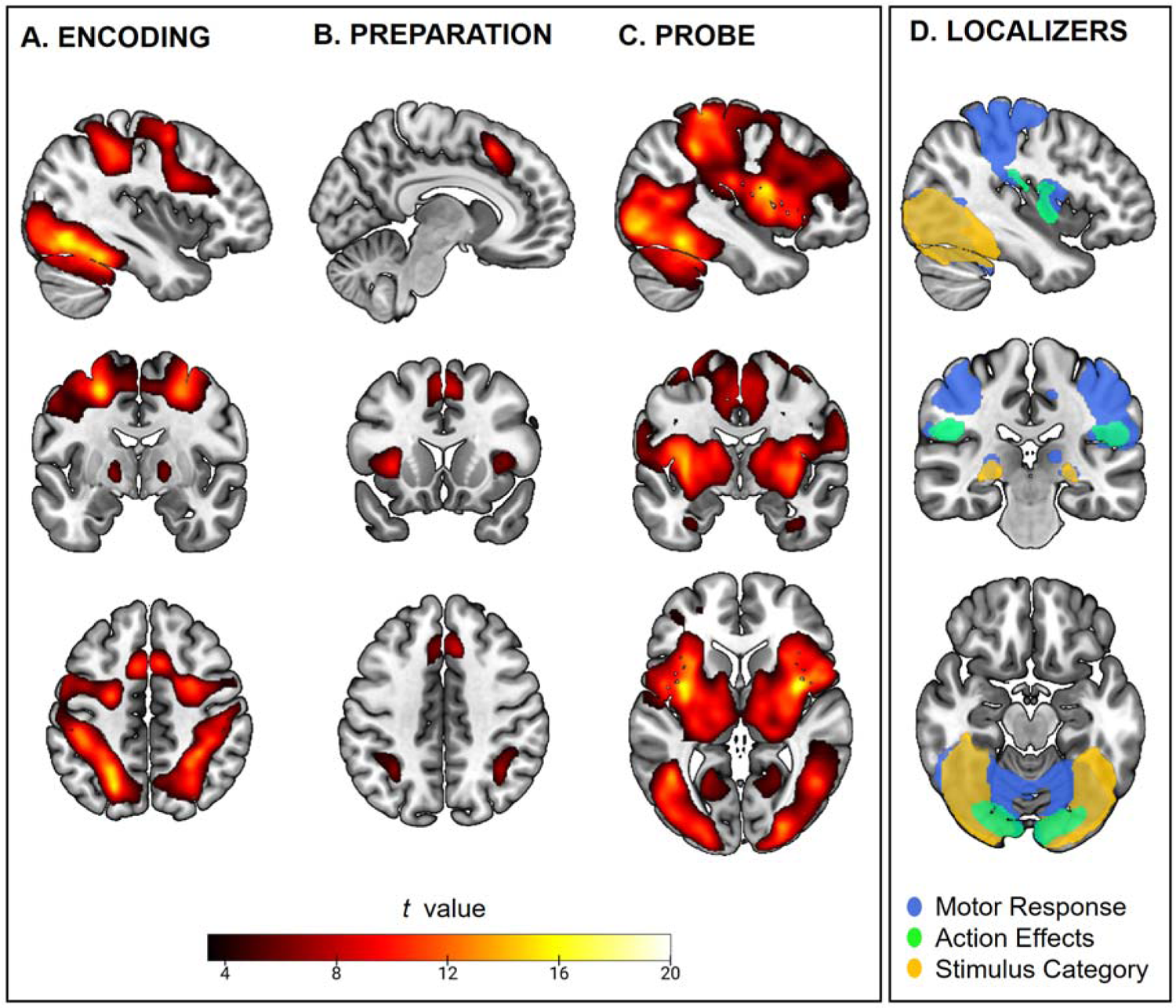
Univariate analysis results. **A-C.** *t* maps showing the increases in activity associated with the encoding (A), preparation (B), and implementation (C) of the novel S-R mappings. **D.** *t* maps showing the increases in activity associated with the three localizers’ event of interest: the response in the motor response localizer (in blue), the tactile stimulation in the action effects localizer (in green), and the animate picture visualization in the stimulus category localizer (yellow). All maps are statistically thresholded at *p* < .05. The unthresholded whole-brain maps used to generate this figure are available in NeuroVault: https://neurovault.org/collections/XQFLCDUH/

With a similar approach, we described the regions engaged by the three localizers’ events of interest (Fig. 4D). As expected, the motor response localizer induced an increase in activity in regions associated with action control. A wide cluster ([20, −88, −12], k = 32523, T(1,29) = 10.60, *p* < .001) bilaterally covered the motor cortices, including M1, premotor cortices, and SMA, and extended into the S1, SPL, and IPL. As expected from the presence of visual stimulation during this localizer, the cluster also included early and late visual cortices. Two additional clusters were found in the basal ganglia and posterior insula in both hemispheres (left: [-46, 0, 12], k = 2418, T(1,29) = 8.45, *p* < .001; right [28, 0, 2], k = 1223, T(1,29) = 7.14, *p* < .001). Conversely, the tactile stimulation applied in the action effect localizer was linked to increased activity in the pInsula, bilaterally (left: [−46, −20, 14, k = 950, T(1,29) = 8.63, *p* < .001; right: [50, −24, 22], k = 1046, T(1,29) = 7.29, *p* < .001).

This task also recruited the early visual cortex ([24, −90, 4], k = 4269, T(1,29) = 13.65, *p* < .001), . Finally, the stimulus category localizer was associated with heightened activity in a cluster covering the bilateral early and late visual cortex and the right SPL and IPL (left: [−38, −84, −2], k = 7009, T(1,29) = 14.43, *p* < .001; right: [40, −82, −4], k = 8761, T(1,29) = 15.52, *p* < .001), and another cluster located in the left SPL and IPL ([−22, −48, 40], k = 575, T(1,29) =6.54, *p* < .001). Increased activity was also found in the bilateral hippocampus (left: [-22, 30, 0], k = 224, T(1,29) = 7.26, *p* < .001; right: [22, −30, −2], k = 273, T(1,29) = 7.41, *p* < .001).

#### Canonical template tracking: ROI analyses

To address the main goal of this work, we conducted a series of multivariate, ROI-based analyses which informed on whether the preparation to implement novel S-R mappings activated templates of the instructed responses and stimuli. On one hand, we explored how the novel mappings’ responses were represented, using templates estimated from the motor response and the action effects localizers. On the other, we also addressed the anticipatory encoding of the mappings’ stimuli, using templates from the stimulus category localizer.

#### Response-related analyses

The results obtained using the motor response templates are displayed in Fig. 5A and **Supp. Table 1**. First, we assessed whether the relevant motor response templates were activated to a greater degree than the irrelevant ones, using the latter as baseline. The results supported this hypothesis in most of the regions explored. The paired-sample Wilcoxon tests were significant in the left and right M1 and S1, SMA, PMd, IPL, SPL, and pInsula. Additional one-sample tests showed that the relevant templates’ beta coefficients were greater than zero in all these regions and the PMv. Finally, after visually inspecting our data, we observed that some regions’ irrelevant templates were linked to below-zero beta coefficients. The corresponding one-sample tests on irrelevant template data evidenced significant below-zero beta weights in the left and right M1 and S1, and the pInsula. Contrary, significant above-zero irrelevant templates’ beta weights were found in the PMv. As expected, neither test was significant in the control ROI, V1.

**Figure 5.**
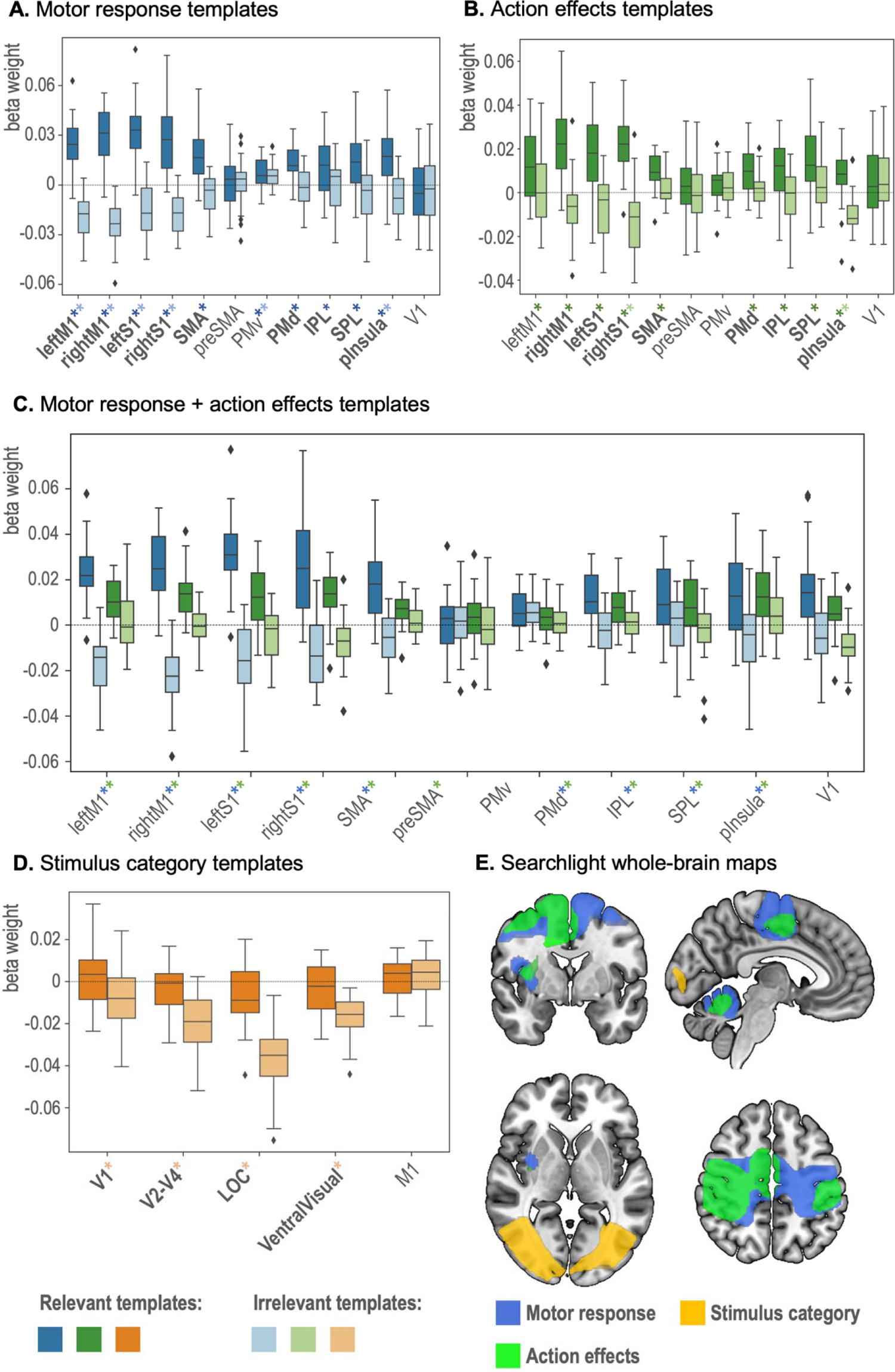
Canonical template tracking results. **A-D:** Boxplots displaying the beta weights for relevant and irrelevant canonical templates. **A.** Motor response localizer results. **B.** Action effects localizer results. **C.** Results after including the motor response and the action effect templates in the same multiple regressions. **D.** Stimulus category localizer results. In **A**, **B**, and **D**, bold ROI labels indicate that the relevant beta weights are significantly greater than the irrelevant ones, and the asterisks, that either the relevant or the irrelevant beta weights (shown in the corresponding darker or lighter color shade) are significantly different than zero. In **C,** the asterisks indicate that, for a given localizer (indicated by the asterisk color, blue: motor response, green: action effects), the relevant beta weights are significantly greater than the irrelevant ones. The statistics (z value, uncorrected and corrected p values) from the corresponding Wilcoxon Signed-Ranked tests are available in **extended** figures 5-1 (results shown in **A** and **B**), **5-2** (results shown in **C**) and **5-3** (results shown in **D**). Average relevant and irrelevant beta weights across individual response and category conditions are displayed in **extended** figures 5-4 (motor response templates), **5-5** (action effects templates), and **5-6** (stimulus category templates). **E.** Searchlight whole-brain maps, showing clusters with significantly higher beta weights from relevant than irrelevant templates. The unthresholded whole-brain maps used to generate this figure are available in NeuroVault: https://neurovault.org/collections/XQFLCDUH/

Next, we applied the same analytical approach with the templates obtained from the action effects localizer (Fig. 5B, **Supp. Table 1**). Like the motor response, the beta weights from relevant and irrelevant action effect templates differed in most of the ROIs. The effect was significant in the right M1, the left and right S1, the SMA, PMd, IPL, SPL, and pInsula. The relevant templates were linked to statistically significant above-zero beta weights across all these regions and the left M1. Finally, we found below-zero beta coefficients linked to irrelevant action effects templates in the left S1 and pInsula. Again, neither test was significant in V1.

Considering the equivalent pattern of results obtained with the two response-related localizers and the nature of both tasks, our findings could also reflect that the motor response and action effects templates captured overlapping representations, and consequently, shared variance from the mappings’ activity patterns. To estimate the unique variance explained by each localizer, we ran an additional analysis including the templates from both tasks simultaneously in the multiple regressions. The resulting averaged beta coefficients are displayed in Fig. 5C **and Supp. Table 2**. To explore these results, we ran repeated measures ANOVA on the beta weights, using ROI (left M1, right M1, left S1, right S1, SMA, preSMA, PMv, PMd, IPL, SPL pInsula), Localizer (motor response, action effects), and Template (relevant, irrelevant) as factors. We found significant main effects of Template, F(1, 26) = 216.65, *p* < .001, η²_p_ = .893, and ROI, F(5.80, 150,80) = 2.76, *p* = .015, η²_p_ = .096, but not of Localizer, F(1, 26) = 0.25, *p* = .619, η²_p_ = .010. Nonetheless, the interaction between Localizer and Template was significant, F(1, 26) = 19.60, *p* < .001, η²_p_ = .430, supporting that the localizer task modulated the relationship between relevant and irrelevant templates. To address the anatomical extent of this effect, we next focused on the significant three-way Localizer x Template x ROI interaction, F(4.89, 127.07) = 18.20, *p* < .001, η²_p_ = .412, and ran post hoc comparisons contrasting relevant and irrelevant templates separately for each localizer and ROI (**Supp. Table 2**). These tests showed a generalized increase in beta coefficients in relevant than irrelevant templates in the two localizer tasks. Significant differences involving motor response and action effect templates were found in left and right M1 and S1, SMA, PMd, IPL, SP, and pInsula. The test was also significant for the action effect templates in the preSMA. Overall, even when the templates’ effect was numerically different depending on the localizer (as evidenced by the two-way and three-way significant interactions), it persisted after including the two localizers’ data simultaneously in the analysis, suggesting that both sets of templates captured independent aspects of the S-R mapping task data.

#### Response templates: exploratory analyses

In addition to our planned analyses, we further explored the response attributes that the motor response and action effects templates were capturing. The visual inspection of the motor response (**Supp.** Fig. 1) and action effect (**Supp.** Fig. 2) beta weights across the six response conditions highlighted the robust effect of response laterality, especially, in primary and secondary somatomotor areas. That could be identified in trials where the division between relevant and irrelevant templates was mapped into different laterality (i.e., in the mappings’ response conditions [left index, left middle] and [right index, right middle]). To rule out that our results were uniquely driven by differences between right- and left-hand response representations, we first compared relevant and irrelevant templates in a subset of trials in which only the effector information was relevant for the analysis. The results from the motor response localizer (Fig. 6A) showed greater beta weights for relevant than irrelevant templates in the right S1 (*z* = 2.76, *p* = .002, *p_corrected_*= .032). A similar pattern was found in the right M1 (*z* = 2.24, *p* = .013, *p_corrected_*= .124) and the SPL (*z* = 1.71, *p* = .044, *p_corrected_* = .394), although these tests did not survive the correction for multiple comparisons. Regarding the action effects templates (Fig. 6B), we found a significant effect in the left S1 (*z* = 2.80, *p* = .003, *p_corrected_*= .028). Next, we aimed to shed some light on the representations instantiated in M1 and S1. Since these two regions have clear contralateral neural architecture, we analyzed separately the two hemispheres’ data. Specifically, we used data from the contralateral hemisphere of specific mappings and templates to perform the analysis. The tests contrasting relevant and irrelevant templates remained significant in S1 when using data from either the motor response (Fig. 6C) or the action effect localizers (Fig. 6D). Specifically, we found a significant effect of the motor response templates in the right S1 (*z* = 2.67, *p* = .004, *p_corrected_* = .016), and the action effects templates in the left S1 (*z* = 2.97, *p* = .002, *p_corrected_* = .002). An equivalent pattern was found in the left and right M1, although neither test survived the statistical correction (Motor response, right M1: *z* = 1.60, *p* = .054, *p_corrected_*= .188; Action effects, left M1: *z* = 1.50, *p* = .066, *p_corrected_*= .081).

**Figure 6.**
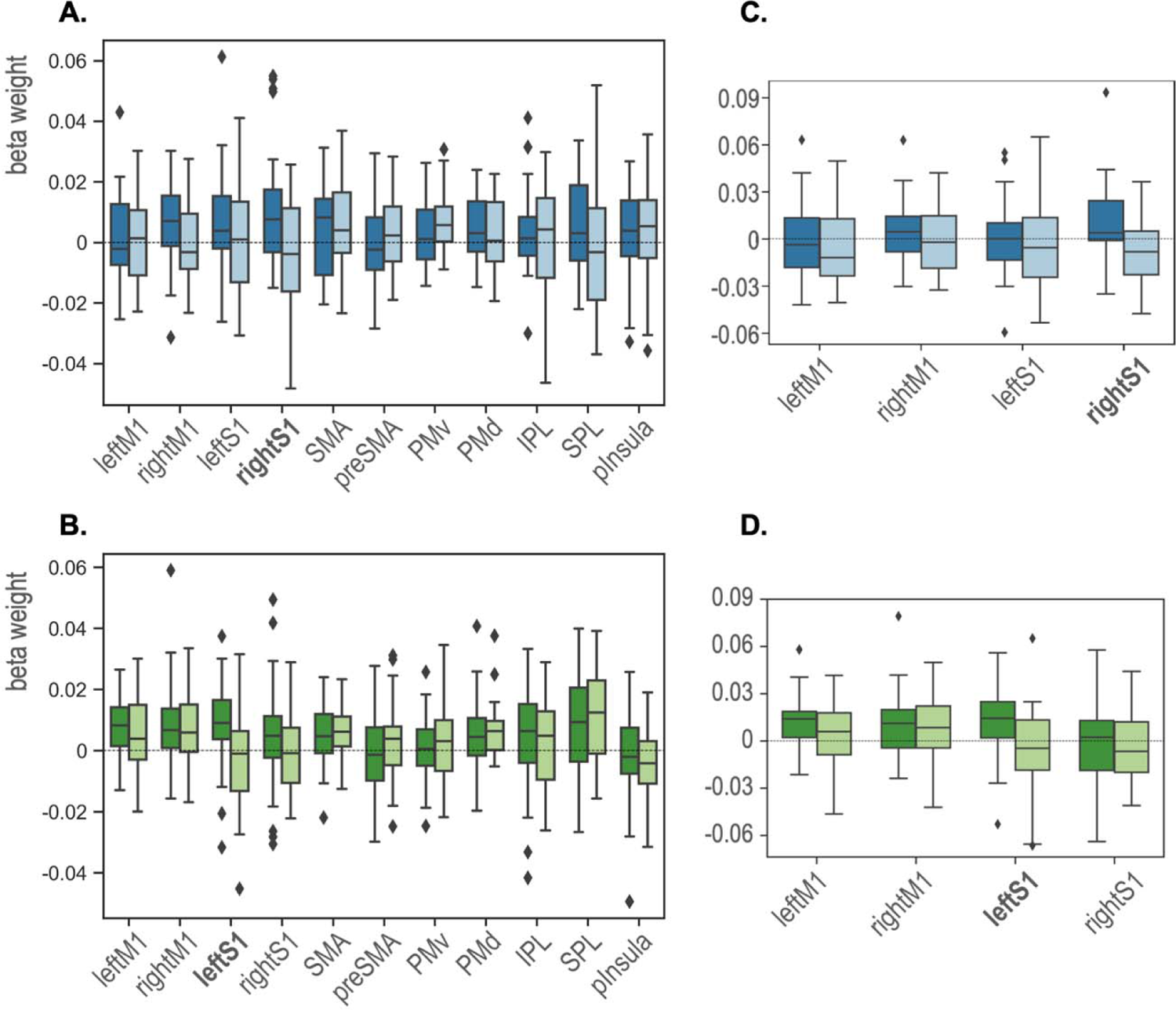
Boxplots showing the beta weights for relevant and irrelevant templates capturing the mappings’ response effector information. **A,B.** Results from the [Left Index, Right Index] and [Left Middle, Right Middle] trials, using the motor response (**A**) and action effects (**B**) templates. **C,D.** Results using activity patterns from the M1 and S1 ROIs contralateral to the mappings’ and canonical templates’ laterality condition, using data from the motor response (**C**) and action effects (**D**) localizer. The ROI labels shown in bold indicate that the relevant beta weights are significantly greater than the irrelevant ones after controlling for multiple comparisons.

Finally, we carried out a series of analyses restricted to the mapping responses that were prepared but not implemented, to ensure that our main results were not driven by motoric signals spreading from the task execution epoch. We repeated the ROI-based canonical template procedure including, in the multiple regression, a single relevant template corresponding to the non-executed mapping response. In general terms, these results replicated most of the reported effects. Using the motor response templates, the paired-sample tests comparing relevant irrelevant templates were significant in left M1 (*z* = 4.38, *p* < .001, *p_corrected_* < .001), right M1 (*z* = 4.75, *p* < .001, *p_corrected_* < .001), left S1 (*z* = 4.22, *p* < .001, *p_corrected_* < .001), right S1 (*z* = 4.53, *p* < .001, *p_corrected_* < .001), SMA (*z* = 4.05, *p* < .001, *p_corrected_* < .001), PMd (*z* = 3.13, *p* < .001, *p_corrected_* = .002), SPL (*z* = 2.37, *p* = .009, *p_corrected_* < .020) and pInsula (*z* = 3.50, *p* < .001, *p_corrected_* < .001). The relevant template beta weights, associated with the prepared but not executed response, were statistically greater than zero in left M1 (*z* = 3.71, *p* < .001, *p_corrected_* < .001), right M1 (*z* = 3.71, *p* < .001, *p_corrected_* < .001), left S1 (*z* = 4.29, *p* < .001, *p_corrected_* < .001), right S1 (*z* = 4.10, *p* < .001, *p_corrected_* < .001) SMA (*z* = 3.51, *p* < .001, *p_corrected_* < .001), PMv (*z* = 3.34, *p* < .001, *p_corrected_* = .003), PMd (*z* = 3.81, *p* < .001, *p_corrected_* < .001), SPL (*z* = 2.44, *p* = .015, *p_corrected_* = .031) and pInsula (*z* = 2.91, *p* = .003, *p_corrected_*= .009). The irrelevant templates were associated with statistically significant below-zero beta weights only in the left M1 (*z* = −3.01, *p* = .003, *p_corrected_* = .014).

Equivalent tests performed with the action effects templates showed greater relevant than irrelevant beta weights in right M1 (*z* = 3.28, *p* < .001, *p_corrected_*= .004), left S1 (*z* = 3.30, *p* < .001, *p_corrected_* = .004), right S1 (*z* = 3.54, *p* < .001, *p_corrected_*< .001), PMd (*z* = 2.46, *p* = .007, *p_corrected_* = .034) and pInsula (*z* = 3.26, *p* < .001, *p_corrected_* = .007). The relevant template beta weights were greater than zero in left M1 (*z* = 2.93, *p* = .003, *p_corrected_*= .011). right M1 (*z* = 3.77, *p* < .001, *p_corrected_*< .001), left S1 (*z* = 3.03, *p* = .002, *p_corrected_*= .008), right S1 (*z* = 3.53, *p* < .001, *p_corrected_*< .001), SMA (*z* = 3.24, *p* = .001, *p_corrected_*= .004), PMd (*z* = 3.48, *p* < .001, *p_corrected_* = .001), and SPL (*z* = 2.98, *p* = .002, *p_corrected_* = .011). Conversely, no significant below-zero beta coefficients were found for the irrelevant templates in this analysis.

Finally, when the motor response and action effect templates were simultaneously included in the multiple regression, significant differences between relevant and irrelevant beta were found for both localizers in left S1 (motor response: *z* = 3.98, *p* < .001, *p_corrected_*< .001; action effects: *z* = 3.11, *p* < .001, *p_corrected_* = .013), right S1 (motor response: *z* = 4.24, *p* < .001, *p_corrected_* < .001; action effects: *z* = 2.90, *p* = .002, *p_corrected_*= .019) and pInsula (motor response: *z* = 2.85, *p* = .002, *p_corrected_*= .014; action effects: *z* = 3.06, *p* = .001, *p_corrected_*= .026). A similar result, although marginally significant for the action effects’ templates, was found in right M1 (motor response: *z* = 4.48, *p* < .001, *p_corrected_* < .001; action effects: *z* = 2.46, *p* = .006, *p_corrected_*= .056). Additional significant results were found for the motor response templates in left M1 (*z* = 4.26, *p* < .001, *p_corrected_*< .001), SMA (*z* = 3.69, *p* < .001, *p_corrected_*< .001) and PMd (*z* = 2.85, *p* = .002, *p_corrected_*= .018).

#### Stimulus-related analyses

We further aimed to address whether the canonical templates obtained during the perception of the instructed stimuli were reactivated during the novel mappings’ preparation. To do so, we followed the same procedure as in the response-related analyses, using templates built from the stimulus category localizer data. The results are displayed in Fig. 5D and **Supp. Table 3**. The paired-sample Wilcoxon tests were significant in all the regions explored, i.e., in V1, V2-4, LOC, and VentralVisual. Nonetheless, we did not find above-zero beta coefficients linked to relevant templates in either region. Instead, the effect was driven by below-zero beta weights associated with the irrelevant templates. The latter effect was significant in the V2-4, LOC, and VentralVIsual ROIs. In these analyses, no effects were detected in the control ROI, M1.

Since the behavioral analyses showed that one of the mappings’ stimulus category conditions (the [bird-fish] combination) was linked to generally impoverished performance, we further explored the template tracking findings across conditions to ensure that the unexpected, below-zero irrelevant beta weights were not driven by an outlier pair of categories. Nonetheless, the pattern found in the main results seemed stable across mappings’ conditions and ROIs (**Supp.** Fig. 3), suggesting that these findings were not driven by trials from a particular condition.

#### Whole-brain searchlight

To complement the ROIs’ results, we carried out an exploratory searchlight analysis. That provided whole-brain maps displaying regions where the beta weights were higher for relevant than irrelevant canonical templates. The results, displayed in Fig. 5E, overlapped substantially with the two sets of ROIs selected a priori for this work. When the motor response localizer templates were used, we found a wide, bilateral cluster ([32, −14, 56], k = 27155, T(1,29) = 17.21, *p* < .001) which covered the motor cortices (M1, PMd and SMA), S1, the IPL and SPL, and the posterior insula. The cluster also extended into the precuneus. A second cluster ([0, −62, −20], k = 5390, T(1,29) = 10.92, *p* < .001) covered the cerebellum bilaterally. Regarding the action effects, we found a significant cluster ([44, −22, 58], k = 13826, T(1,26) = 8.29, *p* < .001) covering the SMA and the right M1 and PMd, and extended into the right S1, IPL, SPL and pInsula. A second cluster involved the left S1, IPL and SPL ([-52, −24, 50], k = 1200, T(1,26) = 4.69, *p* = .027). A significant cluster was also found in the cerebellum ([−20, −60, −26], k = 1718, T(1,26) = 5.18, *p* = .01). Finally, the stimulus category whole-brain map included a significant, bilateral cluster ([50, −76, −2], k = 12470, T(1,29) = 15.21, *p* < .001) extending from the early visual cortices into the fusiform gyrus and the inferotemporal cortex, and which also covered part of the LOC.

To formally address the overlap between the canonical template tracking whole-brain maps and the regions that showed a univariate signal increase during the preparation, we performed three additional conjunction tests, one per localizer task. Overall, these results converged in the anatomical independence between the above-described univariate and multivariate preparatory signals. The motor response map overlapped with the univariate results only in a restricted cluster located in the IPL ([40, −44, 45], k = 64). The same IPL cluster was identified when using the action effect brain map ([41, −44, 45], k = 73), where a small cluster was also found in the posterior section of the pre-SMA ([4, 9, 57], k = 21).

In both cases, the conjunction maps clusters covered an infimal proportion of the significant voxels identified with the template tracking procedure (0.24% and 0.56%, respectively). No conjunction clusters were found when using the stimulus category brain map.

#### Control analysis: template reliability

We aimed to ensure that our results were not biased by differences in reliability across the canonical templates. To do so, we computed and compared (using repeated measures ANOVAs) the SNR and the correlationability across the four template conditions from each localizer. This analysis was carried out, separately, for each of the ROIs explored in the main analysis. In general terms, these results supported that these two reliability measurements were equivalent across template conditions (**Supp.** Fig. 4). We only found significant differences regarding the SNR in action effect templates in the IPL, F(3, 78) = 2.85, *p* < .043, η²_p_ = .099, driven by greater SNR in the right index templates than the right middle finger template. The fact that no other significant result was found in the remaining 67 tests (even when no statistical correction for multiple comparisons was applied) strongly suggests that the findings here reported were not contaminated by differences in the templates’ reliability.

## 4. Discussion

In this work, we aimed to characterize the preparatory neural states leading to novel S-R mapping execution. In contrast to previous literature, which has focused on abstract task-level representations in MDN regions, here we addressed the contribution of lower-level stimulus and response coding across the sensorimotor systems. Critically, we aimed to detect activity patterns with a shared encoding format between novel mapping preparation and overt performance. Using canonical template tracking, we isolated the mappings’ relevant sensorimotor activity patterns and estimated their representational strength during task preparation. Our findings showed that the response-related components were indeed pre-activated during novel mapping preparation, entailing the presence of templates that captured both motor responses and their somatosensory consequences. Nonetheless, we obtained less interpretable results regarding the stimulus-related templates, with non-relevant stimulus representations displaying systematically smaller, below-zero similarity estimates, while the representational strength of the relevant categories did not differ from zero. Overall, our findings support that novel instruction preparation anticipates prospective motor and kinesthetic representations while leaving open the role of stimulus neural codes.

Our work revealed that the encoding of response-related templates was a rather widespread phenomenon across regions previously associated with action control (Haggard, 2008; Rizzolatti & Luppino, 2001), detecting this effect in M1, SMA, PMd, S1, IPL, SPL, and the posterior insula. These regions were not engaged in a univariate fashion during the preparation, pointing out that their anticipatory recruitment did not rely on mean signal increases. On the contrary, we could not evidence template instantiation in the only ROI showing preparatory univariate activation (the pre-SMA). This set of frontoparietal regions not only encoded diverse sensorimotor information (motor responses and action) but also seemed to anticipate, simultaneously, pairs of prospective action plans. In this regard, we replicated our main effects when the analyses were restricted to responses that were prepared but not executed. That also ensured that our findings were not contaminated by response-specific activity originated during task execution. Moreover, the whole-brain searchlight procedure confirmed the anatomical specificity of our findings, showing that the response-related template anticipation was majorly restricted to areas that overlapped with our ROIs. The only exception was the cerebellum which, despite not being included among our a-priori ROIs, has been traditionally associated with action control (Sokolov et al., 2017). Finally, further control analyses supported that the different response-related templates were equally reliable across all the regions explored. These findings validated our analytical approach, ensuring that it was not contaminated by differences in templates’ noise or informational content.

Critically, beyond this overall engagement of the action control system, our evidence also highlighted the role of two areas, M1 and S1, that showed numerically larger and more consistent effects. In both, we found robust above-zero representational strengths for the relevant templates, accompanied by below-zero values for irrelevant ones. We further showed that response effector information was retained after removing the impact of laterality, stressing the specificity of representations held in these areas. Overall, our results stressed the relevance of downstream processing, especially in M1 and S1, during higher-level novel task preparation. Furthermore, we evidenced that these regions’ encoding space was shared with sensorimotor execution, offering a deeper insight into their representational format (Kriegeskorte & Kievit, 2013).

Taken together, these findings fit well within the predictions of simulation-based frameworks (Grush, 2004; Jeannerod, 1994, 2001), suggesting that the task-relevant sensorimotor processing was internally anticipated before the execution. Following previous theoretical (Brass et al., 2017; Moran & O’Shea, 2020) and empirical work (Liefooghe et al., 2021; Theeuwes et al., 2018), we could interpret our results as reflecting the involvement of motor imagery when preparing for novel tasks. In this regard, recent behavioral evidence showed that novel task preparation relied on the pre-activation of effector-specific response representations, since blocking their access (via dual-task demands) robustly impaired performance (Palenciano et al., 2021). Our results seem to capture the neural mechanism underpinning that behavioral effect, evidencing the presence of low-level response representations, and further characterizing the different action-related information that could be anticipated via motor simulation. Also supporting this view, the regions that showed anticipatory template instantiation have been linked to motor imagery in the past (Hardwick et al., 2018; Hétu et al., 2013). Moreover, some of them encode movements in a common representational space that generalizes between imagery and overt execution (Zabicki et al., 2017), as we found in our data. Still, this view contrasts with theoretical accounts stressing that imagery requires cognitive processes that go beyond covert action simulation (such as conscious experience, action monitoring, inhibition, etc.; O’Shea & Moran, 2017). In this regard, the nature of our task (which substantially differed from standard imagery-based paradigms), left unaddressed whether the reported sensorimotor pre-activation reflected explicit, conscious action imagery, or instead, a rather implicit simulation process. Future investigation will be key to characterize the cognitive implications of our findings.

Alternatively, our response-related template tracking results could be also interpreted from broader frameworks on action control. There, it has been widely documented how the somatomotor system encodes prepared actions before their execution (e.g., Gallivan, Adam McLean, et al., 2011; Gallivan, McLean, et al., 2011). Critically, and parallelizing our findings, recent data have supported a representational overlap between motor planning and execution in M1 and S1 (Ariani et al., 2020). From this view, our results could reflect that, in general terms, preparatory signals push the primary somatomotor cortices closer to execution-induced states (Churchland et al., 2010; Shenoy et al., 2013), facilitating performance. However, there is a key difference between this literature and our research: while these studies investigated how a single action is planned, we focused on the preparation of pairs of equally probable actions. Hence, our findings could help expand these models, showing how several action plans can be anticipated in parallel, to be later selected in a goal-oriented fashion (Cisek & Kalaska, 2010).

Moreover, our findings also highlighted the role of task-relevant action effects (in this case, their somatosensory consequences) during instruction preparation. This result is highly relevant for theoretical frameworks based on ideomotor principles, emphasizing that action planning is the consequence of pre-activating the intended perceptual effects (Hommel, 2009; Hommel et al., 2001; Shin et al., 2010). Agreeing with this view and previous neuroimaging findings (Waszak et al., 2012), we showed that task-relevant kinesthetic outputs were anticipated before mapping execution. Nonetheless, our work extended this research by identifying finer-grained action effect representations, and more critically, by evidencing this effect for the first time in more volatile, novel task scenarios. These promising results stress the need to integrate the role of somatosensory action contingencies in current models of instructed behavior, and further investigate how upcoming action effects bind with other novel task components (such as the instructed responses or stimuli). Furthermore, our results showed that the motor response and action effects templates captured independent facets of the preparatory activity patterns. This result is less straightforward to interpret from the above-mentioned ideomotor perspective, which would predict a greater overlap between both sets of templates. In opposition, our findings could reflect that information about the motor responses and their kinesthetic consequences may be segregated and multiplexed across these regions’ representational spaces. Alternatively, they could reflect that each set of templates could also contain non-shared response information. For instance, the response templates could capture additional proprioceptive information beyond the tactile feedback. Future research will be key to describing in greater detail the nature of these representations as well as their functional role during preparation.

Contrary to the response-related results, we obtained a less clear pattern regarding the stimulus templates. We found the expected relationship between prepared and non-prepared target categories, with smaller representational strength for the latter than the former. ROI and whole-brain analyses showed that this pattern spread across the visual stream, with numerically greater effects in higher-order areas sensitive to abstract category information, such as the LOC (Grill-Spector et al., 1999). However, while the irrelevant category templates were consistently linked to negative estimates, the relevant ones were not statistically different from zero. While this finding may relate to the nature of visual imagery (which may differ from the motor domain), we believe that this pattern could be driven by the design of the category localizer task. Since participants responded to the broader animate-inanimate distinction, the individual templates’ animal categories remained task-irrelevant. This decision allowed isolating the stimulus processing from unwanted motor confounds, and it was in line with the action-effect localizer, where the effector-specific somatosensory contingencies were also task-irrelevant. Still, in the case of the stimulus category localizer, this design may have hindered the category information conveyed by the templates. Moreover, the higher variability of the four targeted animate categories (which conveyed multiple, unique exemplars) may have furthers impaired the robustness of these templates. Having said that, the consistent results linked to the irrelevant templates suggest that this information was somehow anticipated. Although it could be tempting to interpret this finding as the suppression of irrelevant category representations during preparation, the similarity measurements computed (i.e., non-cross-validated; Walther et al., 2016) require that relative instead of absolute activation estimates are considered (Palenciano et al., 2023; Walther et al., 2016). Hence, no inference should be performed based only on the irrelevant categories’ below-zero representational strength. Instead, we encourage caution when interpreting these results and emphasize that more compelling evidence is required to understand the significance of stimulus-related representations in this context.

Overall, our results highlighted the action-oriented nature of proactive novel task control, and more specifically, the involvement of downstream motor and somatosensory regions. This view could seem in conflict with previous literature which has stressed the role of cognitive control-related MDN areas during novel instruction processing (Cole et al., 2010; Demanet et al., 2016; Dumontheil et al., 2011; Hartstra et al., 2011, 2012; Palenciano, González-García, Arco, & Ruz, 2019; Ruge & Wolfensteller, 2010). Therefore, the main frameworks explaining instructed performance have focused on how these frontoparietal regions flexibly generate task representation from scratch (Brass et al., 2009, 2017; Cole, Laurent, et al., 2013). We also supported this notion, showing heightened activity during mapping preparation across the MDN. Nonetheless, our methodology enabled us to identify a co-existent preparatory mechanism outside this network and anchored in lower-level cortices. We believe that our findings reflect a different but complementary facet of novel task processing that entails the anticipatory tuning of sensorimotor systems via motor and kinesthetic template instantiation. In this regard, our results left open a key question: how do the higher-order task representations in the MDN, and the sensorimotor coding here described, interact? Current working memory models state that abstract, multidimensional representations held in the MDN (especially, in the lateral prefrontal cortex) lead to the activation of more specific content encoded in the sensory regions (Christophel et al., 2017; Curtis & D’Esposito, 2003; Sreenivasan et al., 2014). Following this framework, one could expect that instruction representations previously found in the MDN precede and influence the engagement of the concrete sensorimotor templates here reported. Future studies exploiting temporally resolved brain data will be crucial to address this hypothesis and bridge both sets of findings. That would enrich the current understanding of instruction processing, and more generally, help to build more holistic frameworks of anticipatory task control that integrate higher-order and downstream cognitive processes.

## 5. Conclusions

In this work, we addressed whether novel S-R mapping preparation entailed the pre-activation of sensorimotor representations that overlapped with those guiding task execution. By combining fMRI recording and tailored pattern analyses, we evidenced the anticipatory engagement of templates conveying the mappings’ motor responses and their perceptual consequences. While this effect was widespread across frontoparietal regions linked to action control, it was particularly robust in the primary somatosensory and motor cortices. In contrast, we did not obtain evidence supporting an equivalent pattern regarding stimulus-related perceptual templates. Overall, our findings stress that motoric and kinesthetic representations anchored in the sensorimotor systems are proactively engaged by instructions, suggesting that motor imagery may underpin preparation in novel contexts. More broadly, these results suggest that the anticipatory and flexible tuning of the sensorimotor interface systems could be a key component for current models of cognitive control.

## Credit author statement

AFP: Conceptualization, Methodology, Data collection, Data analysis, Writing - original draft, Writing - review & editing. CG-G: Conceptualization, Methodology, Writing - review & editing. JDH: Conceptualization, Writing - review & editing, Funding acquisition. BL: Conceptualization, Writing - review & editing, Funding acquisition. MB: Conceptualization, Writing - review & editing, Funding acquisition.

## Data and code Accessibility

The raw fMRI BIDS dataset is available in the following OpenNeuro repository: https://openneuro.org/datasets/ds004829. The processed individual and group-level MRI and behavioral data, together with the scripts used for data analyses, are available on OSF: https://osf.io/dq6v5/. Group-level whole-brain maps can be also accessed in the following NeuroVault repository: https://neurovault.org/collections/XQFLCDUH/

## Acknowledgment

This research was supported by grant G00951N of the Flemish Government attributed to BL and JDH. AFP was supported by Grant PAIDI21_00207 of the Andalusian Autonomic Government. CG-G was supported by Grant IJC2019-040208-I and Project PID2020-116342GA-I00 funded by MCIN/AEI/10.13039/501100011033, and Grant RYC2021-033536-I funded by MCIN/AEI/10.13039/501100011033 and by the European Union NextGeneration EU/PRTR. JDH was supported by Methusalem funding from the Special Research Fund (BOF) of Ghent University (reference number: BOF22/MET_V/002). MB was supported by an Einstein Strategic Professorship of the Einstein Foundation Berlin (EPP-2018-483) and by the Deutsche Forschungsgemeinschaft (DFG, German Research Foundation) under Germany’s Excellence Strategy – EXC 2002/1 “Science of Intelligence” (project number: 390523135). BL was supported by the Utrecht University Focus Area on Human-Centered Artificial Intelligence.

