## Supplementary material for "Concurrent response and action effect representations across the somatomotor cortices during novel task preparation": Supp. Fig. or Supp. Table

**Supplementary Table 1**

*Results from the Wilcoxon Signed-Ranked tests using the motor response and action effects templates.*

|  | Motor response localizer | | | | | | | | | Action effects localizer | | | | | | | | |
| --- | --- | --- | --- | --- | --- | --- | --- | --- | --- | --- | --- | --- | --- | --- | --- | --- | --- | --- |
|  | *Relevant vs. irrelevant* | | | *Relevant against 0* | | | *Irrelevant against 0* | | | *Relevant vs. irrelevant* | | | *Relevant against 0* | | | *Irrelevant against 0* | | |
|  | *z* | *p* | *p_corr_* | *z* | *p* | *p_corr_* | *z* | *p* | *p_corr_* | *z* | *p* | *p_corr_* | *z* | *p* | *p_corr_* | *z* | *p* | *p_corr_* |
| leftM1 | 4.78 | <.001 | <.001 | 4.72 | <.001 | <.001 | -4.68 | <.001 | <.001 | 1.93 | .026 | .056 | 3.22 | .001 | .002 | 0.19 | .848 | 1 |
| rightM1 | 4.78 | <.001 | <.001 | 4.70 | <.001 | <.001 | -4.78 | <.001 | <.001 | 4.02 | <.001 | <.001 | 4.40 | <.001 | <.001 | -2.55 | .011 | .188 |
| leftS1 | 4.78 | <.001 | <.001 | 4.76 | <.001 | <.001 | -4.00 | <.001 | <.001 | 3.54 | <.001 | <.001 | 3.63 | <.001 | <.001 | -1.80 | .072 | .201 |
| rightS1 | 4.69 | <.001 | <.001 | 4.72 | <.001 | <.001 | -4.47 | <.001 | <.001 | 4.43 | <.001 | <.001 | 4.47 | <.001 | <.001 | -3.46 | <.001 | .002 |
| SMA | 4.77 | <.001 | <.001 | 4.43 | <.001 | <.001 | -2.05 | .041 | .057 | 3.52 | <.001 | <.001 | 3.92 | <.001 | <.001 | 0.77 | .442 | 1 |
| preSMA | 0.45 | .325 | .678 | 0.57 | .572 | .507 | 0.85 | .393 | 1 | 2.01 | .022 | .056 | 1.27 | .203 | .258 | -0.02 | .980 | 1 |
| PMv | 0.91 | .183 | .678 | 3.40 | <.001 | .001 | 3.75 | <.001 | <.001 | 0.76 | .225 | .210 | 2.23 | .025 | .083 | 1.11 | .269 | 1 |
| PMd | 4.26 | <.001 | <.001 | 4.45 | <.001 | <.001 | -0.69 | .491 | 1 | 3.71 | <.001 | <.001 | 3.92 | <.001 | <.001 | 0.82 | .414 | 1 |
| IPL | 3.17 | <.001 | <.001 | 3.03 | .002 | .002 | -0.24 | .813 | 1 | 3.50 | <.001 | <.001 | 3.00 | .003 | .005 | -0.62 | .532 | 1 |
| SPL | 3.78 | <.001 | <.001 | 3.51 | <.001 | <.001 | -1.55 | .120 | .243 | 2.68 | .003 | .003 | 3.82 | <.001 | <.001 | 1.61 | .107 | .512 |
| pInsula | 4.67 | <.001 | <.001 | 3.90 | <.001 | <.001 | -2.52 | .012 | .037 | 4.36 | <.001 | <.001 | 3.24 | .001 | .006 | -3.46 | <.001 | .002 |
| V1 | -0.82 | .795 | .825 | -1.45 | .147 | .357 | -0.79 | .428 | 1 | -0.28 | .609 | .684 | 1.13 | .258 | .394 | 1.58 | .113 | .686 |

**Supplementary Table 2**

*Results of the posthoc comparisons from the repeated measures ANOVA including the motor response and action effects templates simultaneously.*

|  | *Motor response* | | | *Action effects* | | |
| --- | --- | --- | --- | --- | --- | --- |
|  | *z* | *p* | *p_corr_* | *z* | *p* | *p_corr_* |
| left M1 | 4.53 | <.001 | <.001 | 2.51 | .006 | .011 |
| right M1 | 4.53 | <.001 | <.001 | 4.23 | <.001 | <.001 |
| left S1 | 4.53 | <.001 | <.001 | 3.57 | <.001 | <.001 |
| right S1 | 4.43 | <.001 | <.001 | 4.26 | <.001 | <.001 |
| SMA | 4.53 | <.001 | <.001 | 3.33 | <.001 | <.001 |
| preSMA | 0.80 | .210 | .707 | 2.49 | .006 | .018 |
| PMv | 0.88 | .190 | .707 | 0.80 | .210 | .648 |
| PMd | 4.05 | <.001 | <.001 | 3.40 | <.001 | <.001 |
| IPL | 3.23 | <.001 | <.001 | 3.40 | <.001 | <.001 |
| SPL | 3.59 | <.001 | .001 | 2.97 | .002 | .004 |
| pInsula | 4.43 | <.001 | <.001 | 3.98 | <.001 | <.001 |
| V1 | -2.05 | .775 | .816 | -1.36 | .097 | .157 |

**Supplementary Table 3**

*Results from the Wilcoxon Signed-Ranked tests using the stimulus category templates.*

|  | *Relevant vs. irrelevant* | | | *Relevant against 0* | | | *Irrelevant against 0* | | | |
| --- | --- | --- | --- | --- | --- | --- | --- | --- | --- | --- |
|  | *z* | *p* | *p_corr_* | *z* | *p* | *p_corr_* | | *z* | *p* | *p_corr_* |
| V1 | 3.27 | <.001 | <.001 | 0.75 | .452 | .707 | | -2.73 | .006 | .012 |
| V2-4 | 4.55 | <.001 | <.001 | -1.31 | .192 | 1 | | -4.64 | <.001 | <.001 |
| LOC | 4.77 | <.001 | <.001 | -2.97 | .039 | .673 | | -4.78 | <.001 | <.001 |
| VentralVisual | 4.67 | <.001 | <.001 | *-*1.24 | .213 | 1 | | -4.78 | <.001 | <.001 |
| M1 | -0.88 | .812 | .590 | 1.08 | .280 | .157 | | 1.32 | .185 | .252 |

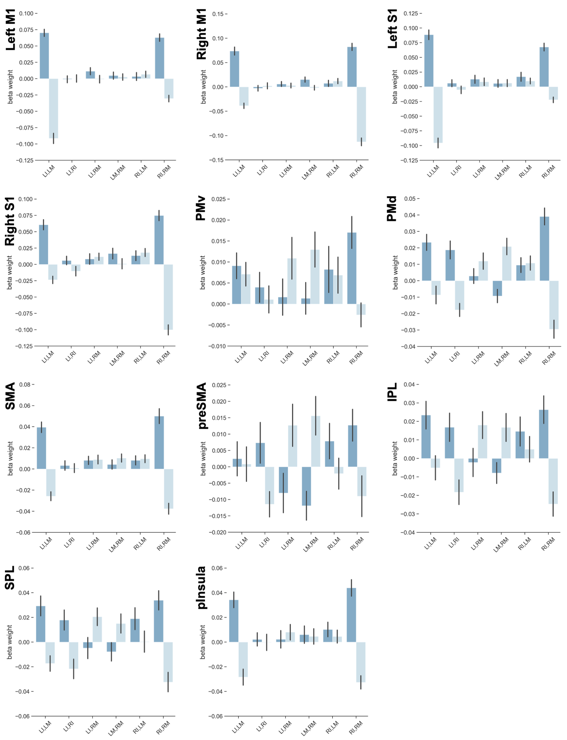

**Supplementary Figure 1.** Bar plots displaying the averaged beta weight for relevant and irrelevant motor response templates, across mappings’ conditions (shown in the X axes) and ROIs. These data were shown for illustrative purposes only. Hence, no statistical information is represented in this figure.

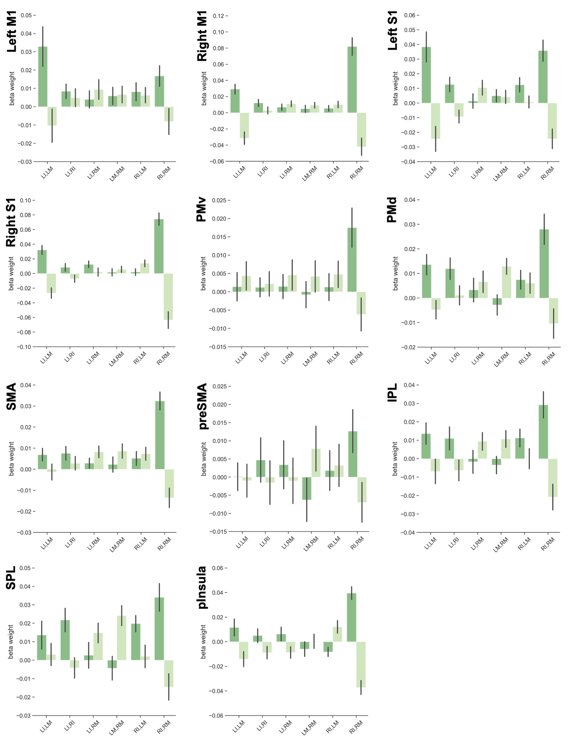

**Supplementary Figure 2.** Bar plots displaying the averaged beta weight for relevant and irrelevant action effects templates, across mappings’ conditions (shown in the X axes) and ROIs. These data were shown for illustrative purposes only. Hence, no statistical information is represented in this figure.

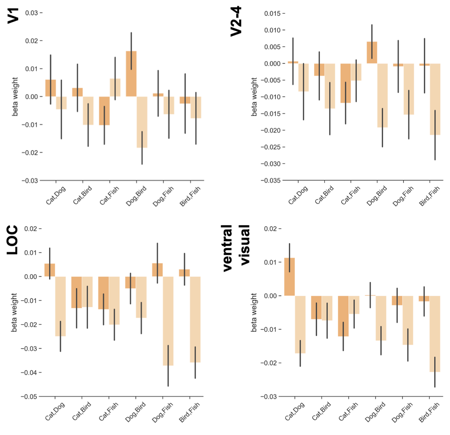

**Supplementary Figure 3.** Bar plots displaying the averaged beta weight for relevant and irrelevant stimulus category templates, across mappings’ conditions (shown in the X axes) and ROIs. These data were shown for illustrative purposes only. Hence, no statistical information is represented in this figure.

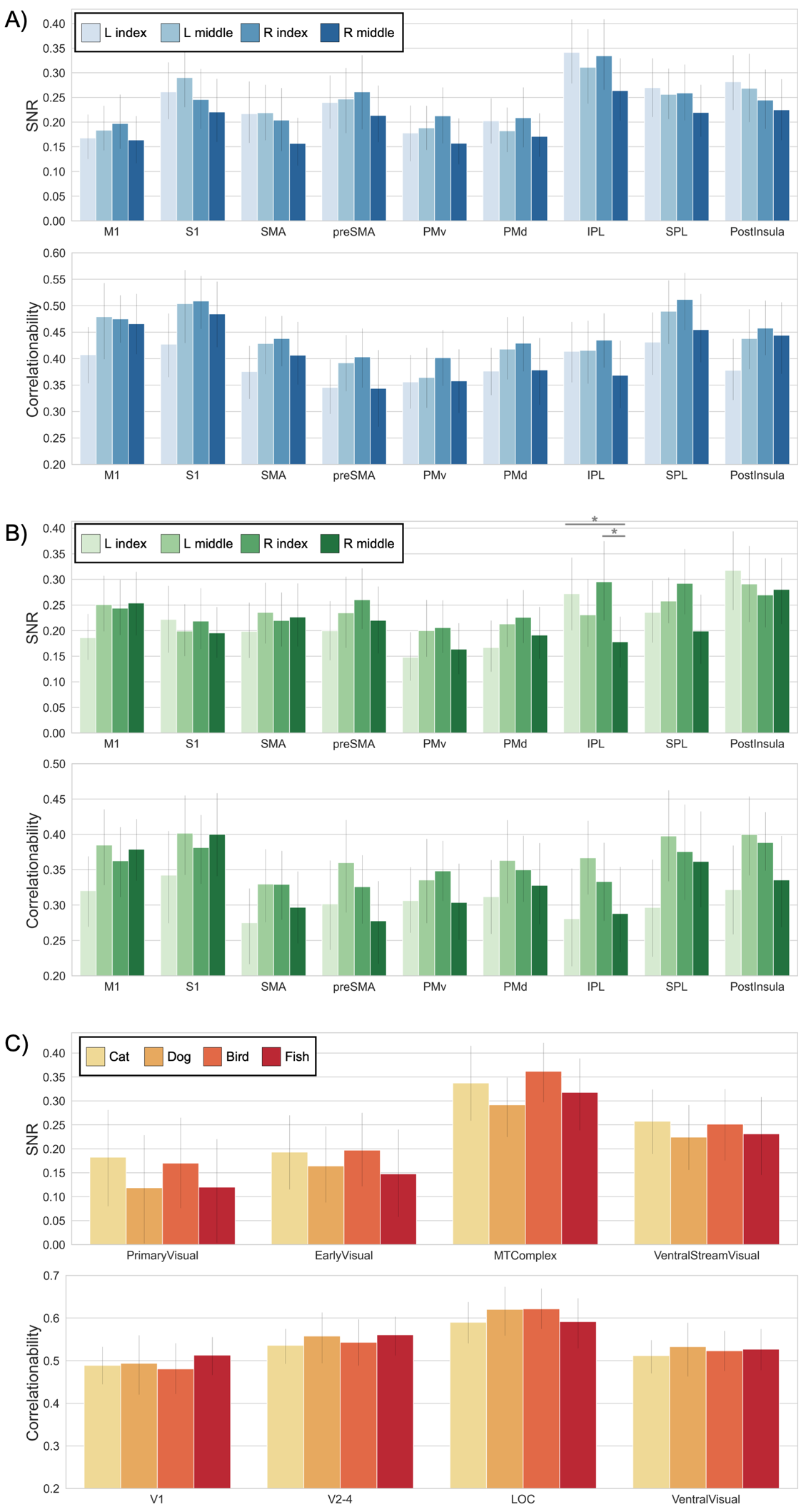

**Supplementary Figure 4**. Canonical templates’ reliability measurements (SNR, correlationability) for the motor response (A), action effects (B), and stimuli category (C) templates, across ROIs. Asterisks denote statistically significant differences (*p* < .05).
